# SWELL1-LRRC8 complex regulates skeletal muscle cell size, intracellular signalling, adiposity and glucose metabolism

**DOI:** 10.1101/2020.06.04.134213

**Authors:** Ashutosh Kumar, Litao Xie, Chau My Ta, Antentor J. Hinton, Susheel K. Gunasekar, Rachel A. Minerath, Karen Shen, Joshua M. Maurer, Chad E. Grueter, E. Dale Abel, Gretchen Meyer, Rajan Sah

## Abstract

Maintenance of skeletal muscle is beneficial in obesity and Type 2 diabetes. Mechanical stimulation can regulate skeletal muscle differentiation, growth and metabolism, however the molecular mechanosensor remains unknown. Here, we show that SWELL1 (LRRC8a) functionally encodes a swell-activated anion channel that regulates PI3K-AKT, ERK1/2, mTOR signaling, muscle differentiation, myoblast fusion, cellular oxygen consumption, and glycolysis in skeletal muscle cells. SWELL1 over-expression in SWELL1 KO myotubes boosts PI3K-AKT-mTOR signaling to supra-normal levels and fully rescues myotube formation. Skeletal muscle targeted SWELL1 KO mice have smaller myofibers, generate less force *ex vivo*, and exhibit reduced exercise endurance, associated with increased adiposity under basal conditions, and glucose intolerance and insulin resistance when raised on a high-fat diet, compared to WT mice. These results reveal that the SWELL1-LRRC8 complex regulates insulin-PI3K-AKT-mTOR signalling in skeletal muscle to influence skeletal muscle differentiation *in vitro* and skeletal myofiber size, muscle function, adiposity and systemic metabolism *in vivo*.

## Introduction

Maintenance of skeletal muscle mass and function is associated with improved metabolic health and is thought to protect against obesity and obesity-related diseases such as diabetes, nonalcoholic fatty liver disease, heart disease and osteoarthritis, and correlates with overall health in the aging population. Skeletal muscle atrophy, the loss of skeletal muscle mass, is associated with cancer (cachexia), heart failure, chronic corticosteroid use, paralysis or denervation (disuse atrophy), chronic positive pressure ventilation (diaphragm atrophy, inability to extubate), prolonged space flight^1^ or bed rest (unloading) and aging^2^. Each of these clinical scenarios also contribute to poor metabolic health and increase frailty. Thus, a deeper understanding of the molecular mechanisms that regulate skeletal muscle maintenance, growth and function has important implications for human health.

Skeletal muscle growth is regulated by a multitude of stimuli, including growth factor signaling, cytokines, mechanical load, integrin signaling and hormones^2^. Activation of AKT-mTOR signaling downstream of insulin and IGF-1 is well established as a critical regulator of skeletal muscle differentiation and growth^2^. However, mechanical loading of muscle, as occurs with regular activity, exercise, and resistance training also induces mTOR-mediated skeletal muscle growth^3–5^, possibly via a growth-factor independent mechanosensory mechanism. β1-integrin and focal adhesion kinase (FAK) signaling has been proposed as this mechanosensory system - sensing mechanical load on skeletal muscle and regulating hypertrophic signaling^6–9^. However, mechanosensitive ion channels, or ion channel complexes, may also regulate intracellular signaling, including PIEZO1, TRPV4 and the recently identified SWELL1 (LRRC8a) channel complex that functionally encodes the volume-regulated anion current (VRAC)^10–12^.

*SWELL1* or *LRRC8a* (Leucine-Rich Repeat Containing Protein 8A) encodes a ~95 kDa membrane protein with four transmembrane domains and an intracellular, C-terminal Leucine-Rich Repeat Domain (LRRD)^13^. It was first described, in 2003, as the site of a translocation mutation in a young woman with an immunodeficiency characterized by agammaglobulinemia and absent B-cells^14,15^. This phenotype led to studies linking SWELL1 to lymphocyte differentiation defects arising from impaired SWELL1 dependent GRB2-mediated PI3K-AKT signaling, in part based on data from a global SWELL1/LRRC8a knock-out (KO) mouse^16^. Although SWELL1 was speculated in 2012 to form a hetero-hexameric ion channel complex with other LRRC8 family members^17^, it was not until 2014 that SWELL1/LRRC8a was experimentally identified as an essential component of the volume-regulated anion channel (VRAC)^11,12^, forming hetero-hexamers with LRRC8b-e^10,11^. So, for about a decade, SWELL1 was considered a membrane protein that regulates PI3K-AKT mediated lymphocyte function^14,15^, putatively via non-ion channel, protein-protein interaction mediated signaling, and only later discovered to also form the long-studied VRAC ion channel signaling complex. Indeed, since its first description >30 years ago^18–20^, VRAC has been associated with a multitude of complex physiological and pathophysiological functions, including cell proliferation, cell migration, angiogenesis, cell death and apoptosis^21,22^; however, the molecular mechanisms underlying these diverse functions had remained elusive without knowledge of the molecular identity of this ion channel complex.

We recently identified SWELL1-LRRC8 as a swell or stretch-activated volume sensor in adipocytes that regulates glucose uptake, lipid content, and adipocyte growth via a SWELL1-GRB2-PI3K-AKT signaling pathway – providing a putative feed-forward amplifier to enhance adipocyte growth and insulin signaling during caloric excess^23,24^. Intriguingly, others have shown that mechanical stimuli applied by pulling on β1-integrins on cardiac muscle cells with magnetic beads activates a SWELL1-LRRC8 like current (VRAC), suggesting a putative connection between β1-integrin/focal adhesion kinase signaling and SWELL1^25,26^. Taken together, these findings suggest that SWELL1 may connect β1-integrin mediated mechano-transduction with insulin/IGF1-PI3K-AKT-mTOR signaling, which, in skeletal muscle is anticipated to regulate skeletal muscle differentiation, function and potentially also adiposity and systemic glucose metabolism^27–29^. In this study, we tested this hypothesis by examining SWELL1-LRRC8 dependent intracellular signaling and myotube differentiation in both C2C12 and primary skeletal muscle cells *in vitro*, performing skeletal muscle targeted SWELL1 loss-of-function experiments *in vivo*. We find that myotube differentiation and insulin and stretch-mediated PI3K-AKT, ERK1/2, mTOR signaling is strongly regulated by SWELL1 protein expression, and provide evidence that GRB2 signaling mediates these SWELL1 dependent effects. Finally, using skeletal muscle SWELL1 knock-out mice, we reveal the requirement of skeletal muscle SWELL1 for maintaining normal skeletal muscle cell size, muscle endurance, force generation, adiposity and glucose tolerance, under basal conditions and in the setting of overnutrition.

## Results

### SWELL1 is expressed and functional in skeletal muscle and is required for myotube formation

SWELL1 (LRRC8a) is the essential component of a hexameric ion channel signaling complex that encodes I_CI,SWELL_, or the volume-regulated anion current (VRAC)^11,12^. While the SWELL1-LRRC8 complex has been shown to regulate cellular volume in response to application of non-physiological hypotonic extracellular solutions, the physiological function(s) of this ubiquitously expressed ion channel signaling complex remain unknown. To determine the function of the SWELL1-LRRC8 channel complex in skeletal muscle, we genetically deleted *SWELL1* from C2C12 mouse myoblasts using CRISPR/cas9 mediated gene editing as described previously ^23,30^, and from primary skeletal muscle cells isolated from *SWELL1^flfl^* mice transduced with adenoviral Cre-mCherry (KO) or mCherry alone (WT control)^23^. SWELL1 protein Western blots confirmed robust SWELL1 ablation in both SWELL1 KO C2C212 myotubes and SWELL1 KO primary skeletal myotubes (**Figure 1A**). Next, whole-cell patch clamp revealed that the hypotonically-activated (210 mOsm) outwardly rectifying current present in WT C2C12 myoblasts is abolished in SWELL1 KO C2C12 myoblasts (**Figure 1B**), confirming SWELL1 as also required for I_CI,SWELL_ or VRAC in skeletal muscle myoblasts. Remarkably, SWELL1 ablation is associated with impaired myotube formation in both C2C12 myoblasts and in primary skeletal satellite cells (**Figure 1C**), with an 58% and 45% reduction in myotube area in C2C12 and skeletal muscle myotubes, respectively, compared to WT. As an alternative metric, myoblast fusion is also markedly reduced by 80% in SWELL1 KO C2C12 compared to WT, as assessed by myotube fusion index (number of nuclei inside myotubes/ total number of nuclei; **Figure 1C**).

**Figure 1:**
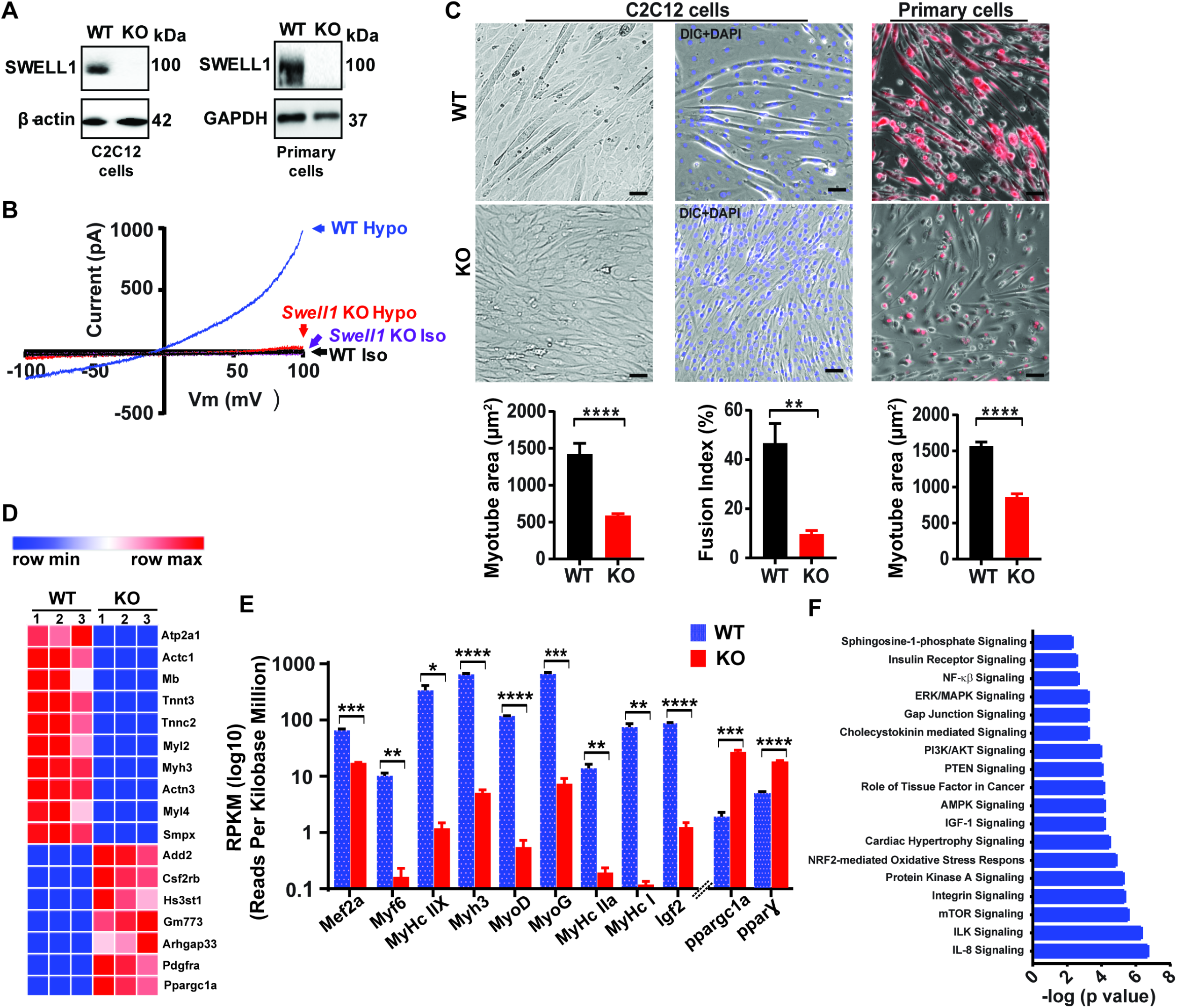
Skeletal muscle SWELL1 is required for myotube formation and regulates multiple myogenic signaling pathways. **A**, Western blots from WT and SWELL1 KO C2C12 (left) and primary myotubes (right). **B**, Current-voltage curves from WT and SWELL1 KO C2C12 myoblast measured during a voltage-ramp from −100 to +100 mV +/- isotonic and hypotonic (210 mOsm) solution. **C**, Bright field merged with fluorescence images of differentiated WT and SWELL1 KO C2C12 myotubes (left, middle) and skeletal muscle primary cells (right). DAPI stains nuclei blue (middle). Red is mCherry reporter fluorescence from adenoviral transduction. Scale bar: 100 μm. Mean myotube surface area measured from WT (n = 21) and SWELL1 KO (n=21) C2C12 myotubes (left), and WT (n = 22) and SWELL1 KO (n = 15) primary skeletal myotubes (right). Fusion index (%multinucleated cells) measured from WT (n = 5 fields) and SWELL1 KO (n = 5 fields) C2C12 (shown below the representative image). **D**, Heatmap of top 17 differentially expressed genes in WT versus SWELL1 KO C2C12 myotubes derived from RNA sequencing. **E**, Reads Per Kilobase Million for select myogenic differentiation genes (n= 3, each). **F**, IPA canonical pathway analysis of genes significantly regulated in SWELL1 KO C2C12 myotubes in comparison to WT. n = 3 for each group. For analysis with IPA, FPKM cutoffs of 1.5, fold change of ≥1.5, and false discovery rate < 0.05 were utilized for significantly differentially regulated genes. Statistical significance between the indicated values were calculated using a two-tailed Student’s t-test. Error bars represent mean ± s.e.m. *, P < 0.05, **, P < 0.01, ***, P < 0.001, ****, P < 0.0001. n = 3, independent experiments.

### Global transcriptome analysis reveals that SWELL1 ablation blocks myogenic differentiation and dysregulates multiple myogenic signaling pathways

In order to further characterize the observed SWELL1 dependent impairment in myotube formation in C2C12 and primary muscle cells we performed genome-wide RNA sequencing (RNA-seq) of SWELL1 KO C2C12 relative to control WT C2C12 myotubes. These transcriptomic data revealed clear differences in the global transcriptional profile between WT and SWELL1 KO C2C12 myotubes (**Figure 1D** and **Supplementary file 1**), with marked suppression of numerous skeletal muscle differentiation genes including Mef2a (0.2-fold), Myl2 (0.008-fold), Myl3 (0.01-fold), Myl4 (0.008-fold), Actc1 (0.005-fold), Tnnc2 (0.005-fold), Igf2 (0.01-fold) (**Figure 1E**). Curiously, this suppression of myogenic differentiation is associated with marked induction of ppargc1α (PGC1α; 14-fold) and PPARƔ (3.7-fold). PGC1α and PPARƔ are positive regulators of skeletal muscle differentiation^31–33^, suggesting that the SWELL1-dependent defect in skeletal muscle differentiation lies downstream of PGC1α and *PPARƔ*. To further define putative pathway dysregulation underlying SWELL1 mediated disruptions in myogenesis we next performed pathway analysis on the transcriptome data. We find that numerous signaling pathways essential for myogenic differentiation are disrupted, including insulin (2X10^−3^), MAP kinase (5X10^−4^), PI3K-AKT (1X10^−4^), AMPK (6X10^−5^), integrin (3X10^−6^), mTOR (2X10^−6^), integrin linked kinase (4X10^−7^) and IL-8 (1X10^−7^) signaling pathways (**Figure 1F** and **Supplementary file 2**).

### SWELL1 regulates multiple insulin dependent signaling pathways in skeletal myotubes

Guided by the results of the pathway analysis, and the fact that skeletal myogenesis and maturation is regulated by insulin-PI3K-AKT-mTOR-MAPK ^34, 2^ we directly examined a number of insulin-stimulated pathways in WT and SWELL1 KO C2C12 myotubes, including insulin-stimulated AKT2-AS160, FOXO1 and AMPK signaling. Indeed, insulin-stimulated pAKT2, pAS160, pFOXO1 and pAMPK are abrogated in SWELL1 KO myotubes compared to WT C2C12 myotubes (**Figure 2A&C**). Importantly, insulin-AKT-AS160 signaling is also diminished in SWELL1 KO primary skeletal muscle myotubes compared to WT primary myotubes (**Figure 2B&D**), consistent with the observed differentiation block (**Figure 1C**). This confirms that SWELL1-dependent insulin-AKT and downstream signaling is not a feature specific to immortalized C2C12 myotubes, but is also conserved in primary skeletal myotubes. It is also notable that reduction in total AKT2 protein is associated with SWELL1 ablation in both C2C12 and primary skeletal muscle cells, and this is consistent with 3-fold reduction in AKT2 mRNA expression observed in RNA sequencing data (**Figure 2E**). Moreover, transcription of a number of critical insulin signaling and glucose homeostatic genes are suppressed by SWELL1 ablation, including GLUT4 (SLC2A4, 51-fold), FOXO3 (2-fold), FOXO4 (2.8-fold) and FOXO6 (18-fold) (**Figure 2E**). Indeed, FOXO signaling is thought to integrate insulin signaling with glucose metabolism^35,36^ in a number of insulin sensitive tissues. Collectively, these data indicate that impaired SWELL1-dependent insulin-AKT-AS160-FOXO signaling is associated with the observed defect in myogenic differentiation upon SWELL1 ablation in cultured skeletal myotubes, and also predict putative impairments in skeletal muscle glucose metabolism and oxidative metabolism.

**Figure 2:**
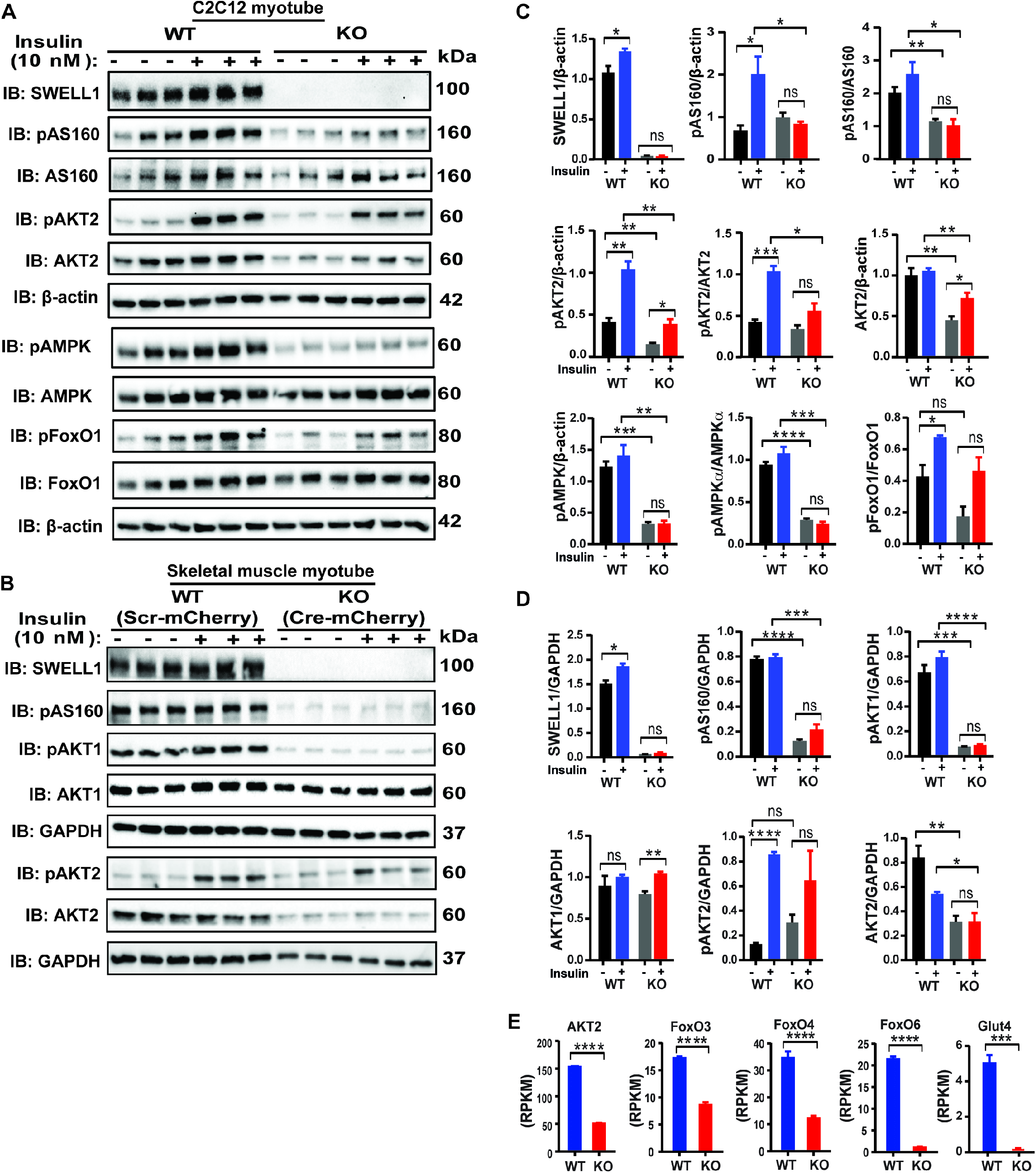
SWELL1 regulates multiple insulin dependent signaling pathways in skeletal myotubes. **A**, Western blots of SWELL1, pAKT2, AKT2, pAS160, AS160, pAMPK, AMPK, pFoxO1, FoxO1 and β-actin in WT and SWELL1 KO C2C12 myotubes upon insulin-stimulation (10 nM). **B**, Western blots of SWELL1, pAS160, pAKT1, AKT1, pAKT2, AKT2 and GAPDH in WT (Ad-CMV-mCherry) and SWELL1 KO (Ad-CMV-Cre-mCherry) primary skeletal muscle myotubes following insulin-stimulation (10 nM). **C**&**D**, Densitometric quantification of proteins depicted on western blots normalized to corresponding β-actin and GAPDH respectively. **E**, Gene expression analysis of insulin signaling associated genes AKT2, FOXO3, FOXO4, FOXO6 and GLUT4 in WT and SWELL1 KO C2C12 myotubes. Statistical significance between the indicated values were calculated using a two-tailed Student’s t-test. Error bars represent mean ± s.e.m. *, P < 0.05, **, P < 0.01, ***, P < 0.001, ****, P < 0.0001. n = 3, independent experiments.

### SWELL1 over-expression in SWELL1 depleted C2C12 is sufficient to rescue myogenic differentiation and augment intracellular signaling above baseline levels

To further validate SWELL1-mediated effects on muscle differentiation and signaling we re-expressed SWELL1 in SWELL1 KO C2C12 myoblasts (SWELL1 O/E) and then examined myotube differentiation and basal activity of multiple intracellular signaling pathways by Western blot, including pAKT1, pAKT2, pAS160, p-p70S6K, pS6K and pERK1/2 as compared to WT and SWELL1 KO C2C12 myotubes. SWELL1 O/E to 2.12-fold WT levels fully rescues myotube development in SWELL1 KO myotubes (**Figure 3A**), as quantified by restoration of SWELL1 KO myotube area to levels above WT (**Figure 3B**). This rescue of SWELL1 KO myotube development upon SWELL1 O/E (**Figure 3A&B**) is associated with either restored (pAS160, AKT2, pAKT1, AKT1, p70S6K) or supra-normal (pAKT2, p-p70S6K, pS6K, pERK1/2) signaling (**Figure 3C&D**) compared to WT C2C12 myotubes. These data demonstrate that SWELL1 protein expression level strongly regulates skeletal muscle insulin signaling and myogenic differentiation.

**Figure 3:**
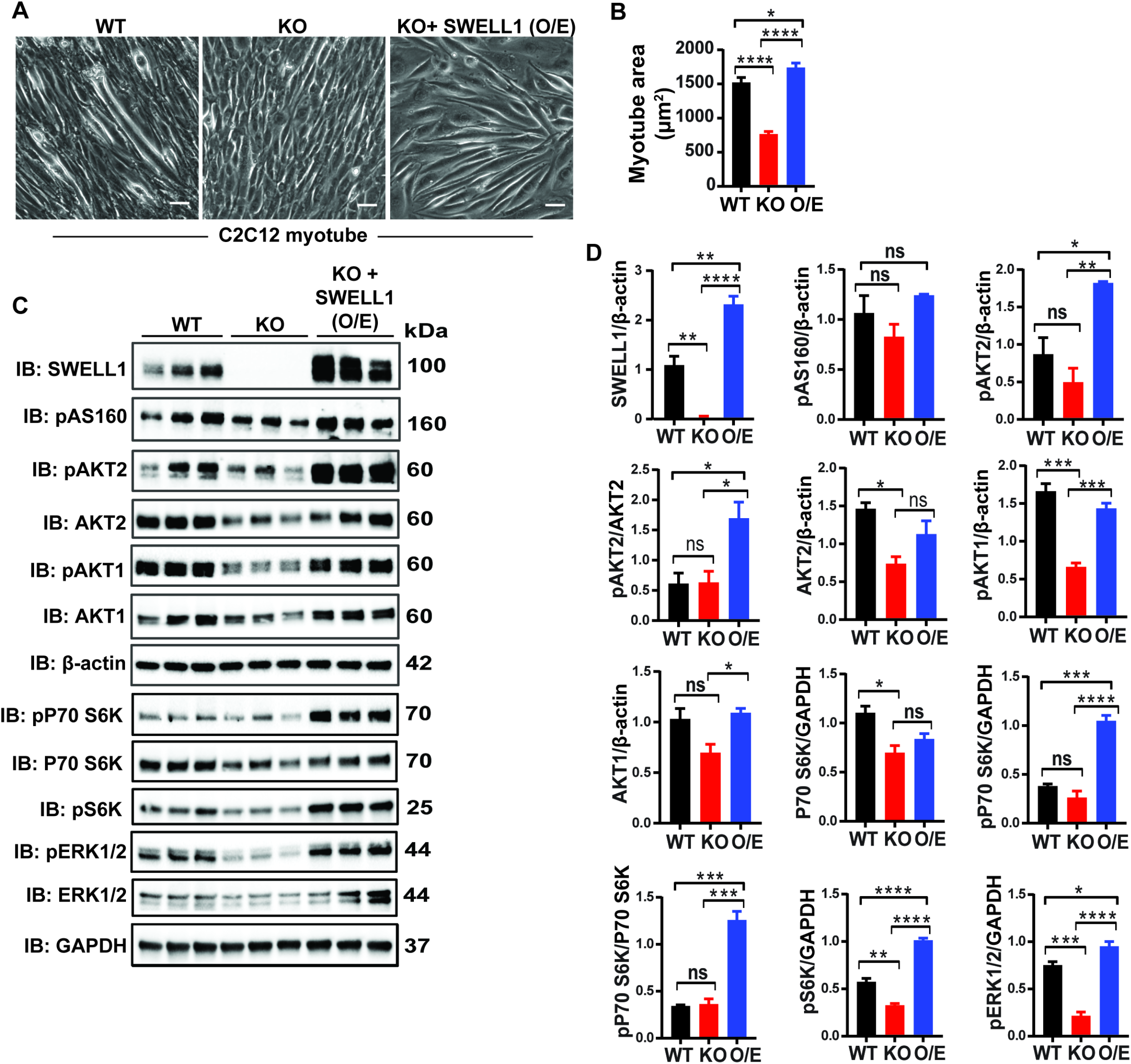
SWELL1 over-expression in SWELL1 KO C2C12 myotubes rescues myogenic differentiation and augments intracellular signaling. **A**, Bright-field image of differentiated WT, SWELL1 KO and SWELL1 KO + SWELL1 O/E C2C12 myotubes. **A**, Quantification of mean myotube surface areas in WT (n = 35), SWELL1 KO C2C12 (n = 26) and SWELL1 KO + SWELL1 O/E C2C12 (n = 45) cells. Scale bar: 100 μm. **C**, Western blots of SWELL1, AKT2, pAKT2, pAS160, pAKT1, AKT1, pP70S6K, P70S6K, pS6K, pERK1/2, ERK1/2, β-actin and GAPDH from WT, SWELL1 KO and SWELL1 KO + SWELL1 O/E C2C12 myotubes. **D**, Densitometric quantification of proteins depicted on western blots normalized to β-actin and GAPDH respectively. Statistical significance between the indicated group were calculated with one-way Anova, Tukey’s multiple comparisons test. Error bars represent mean ± s.e.m. *, P < 0.05, **, P < 0.01, ***, P < 0.001, ****, P < 0.0001. n = 3, independent experiments

### SWELL1-LRRC8 mediates stretch-dependent PI3K-pAKT2-pAS160-MAPK signaling in C2C12 myotubes

In a cellular context, there are numerous reports that VRAC and the SWELL1 -LRRC8 complex that functionally encodes it is mechano-responsive ^25,26 13,37–41^. It is well established that mechanical stretch is an important regulator of skeletal muscle proliferation, differentiation and skeletal muscle hypertrophy and may be mediated by PI3K-AKT-MAPK signaling^2,42,43^ and integrin signaling pathways^6–9^. To determine if SWELL1 is also required for stretch-mediated AKT and MAP kinase signaling in skeletal myotubes we subjected WT and SWELL1 KO C2C12 myotubes to 0% or 5% equiaxial stretch using the FlexCell stretch system. Mechanical stretch (5%) is sufficient to stimulate PI3K-AKT2/AKT1-pAS160-MAPK (ERK1/2) signaling in WT C2C12 in a SWELL1-dependent manner (**Figure 4A&B**). These data position SWELL1-LRRC8 as a co-regulator of both insulin and stretch-mediated PI3K-AKT-pAS160-MAPK signaling.

**Figure 4:**
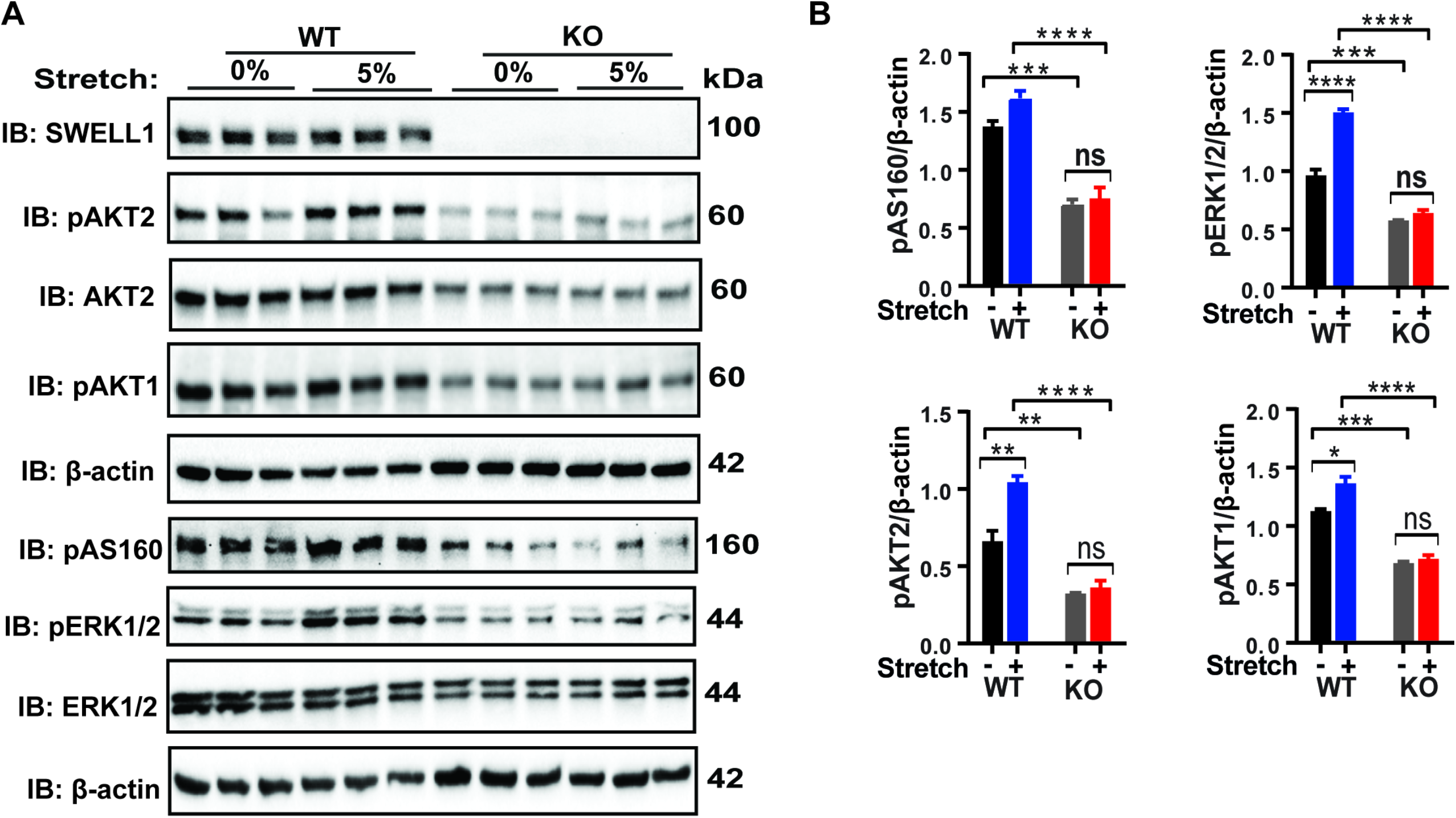
SWELL1 is required for intact stretch-induced PI3K-pAKT2-pAS160-MAPK signaling in C2C12 myotubes. **A**, Western blot of SWELL1, AKT2, pAKT2, pAKT1, pAS160, pERK1/2, ERK1/2 and β-actin in WT and SWELL1 KO myotube in response to 15 minutes of 0% and 5% static stretch. **B**, Densitometric quantification of each signaling protein relative to β-actin. Statistical significance between the indicated group calculated with one-way Anova, Tukey’s multiple comparisons test. Error bars represent mean ± s.e.m. *, P < 0.05, **, P < 0.01, ***, P < 0.001, ****, P < 0.0001. n = 3, independent experiments

### SWELL1 interacts with GRB2 in C2C12 myotubes and regulates myogenic differentiation

It has been reported earlier in both lymphocyte and adipocytes that the SWELL1-LRRC8 complex interacts with Growth factor Receptor-Bound 2 (GRB2) and regulates PI3K-AKT signaling^16,23,24^, whereby GRB2 binds with IRS1/2 and negatively regulates insulin signaling^44^. Indeed, GRB2 knock-down augments insulin-PI3K-MAPK signaling and induces myogenesis and myogenic differentiation genes^44–46^. To determine if SWELL1 and GRB2 interact in C2C12 myotubes, we overexpressed C-terminal 3XFlag tagged SWELL1 in C2C12 cells followed by immunoprecipitation (IP) with Flag antibody. We observed significant GRB2 enrichment upon Flag IP from lysates of SWELL1-3xFlag expressing C2C12 myotubes, consistent with a GRB2-SWELL1 interaction (**Figure 5A**). Based on the notion that SWELL1 titrates GRB2-mediated suppression of AKT/MAPK signaling, and that SWELL1 ablation results in unrestrained GRB2-mediated AKT/MAPK inhibition^24^, we next tested if GRB2 knock-down (KD) may rescue myogenic differentiation in SWELL1 KO C2C12 myotubes. shRNA-mediated GRB2 KD in SWELL1 KO C2C12 myoblasts (SWELL1 KO/shGRB2; **Figure 5B**) stimulates myotube formation (**Figure 5C**) and increases myotube area (**Figure 5D**), to levels equivalent to WT/shSCR (**Figure 5C&D**). Similarly, GRB2 KD in SWELL1 KO C2C12 myotubes induces myogenic differentiation markers IGF1, MyoHCl, MyoHClla and MyoHCIIb relative to both SWELL1 KO/shSCR and WT/shSCR (**Figure 5E&F**). These data are consistent with GRB2 suppression rescuing myotube differentiation in SWELL1 KO C2C12, and supports a model in which SWELL1 regulates myogenic differentiation by titrating GRB2-mediated signaling.

**Figure 5:**
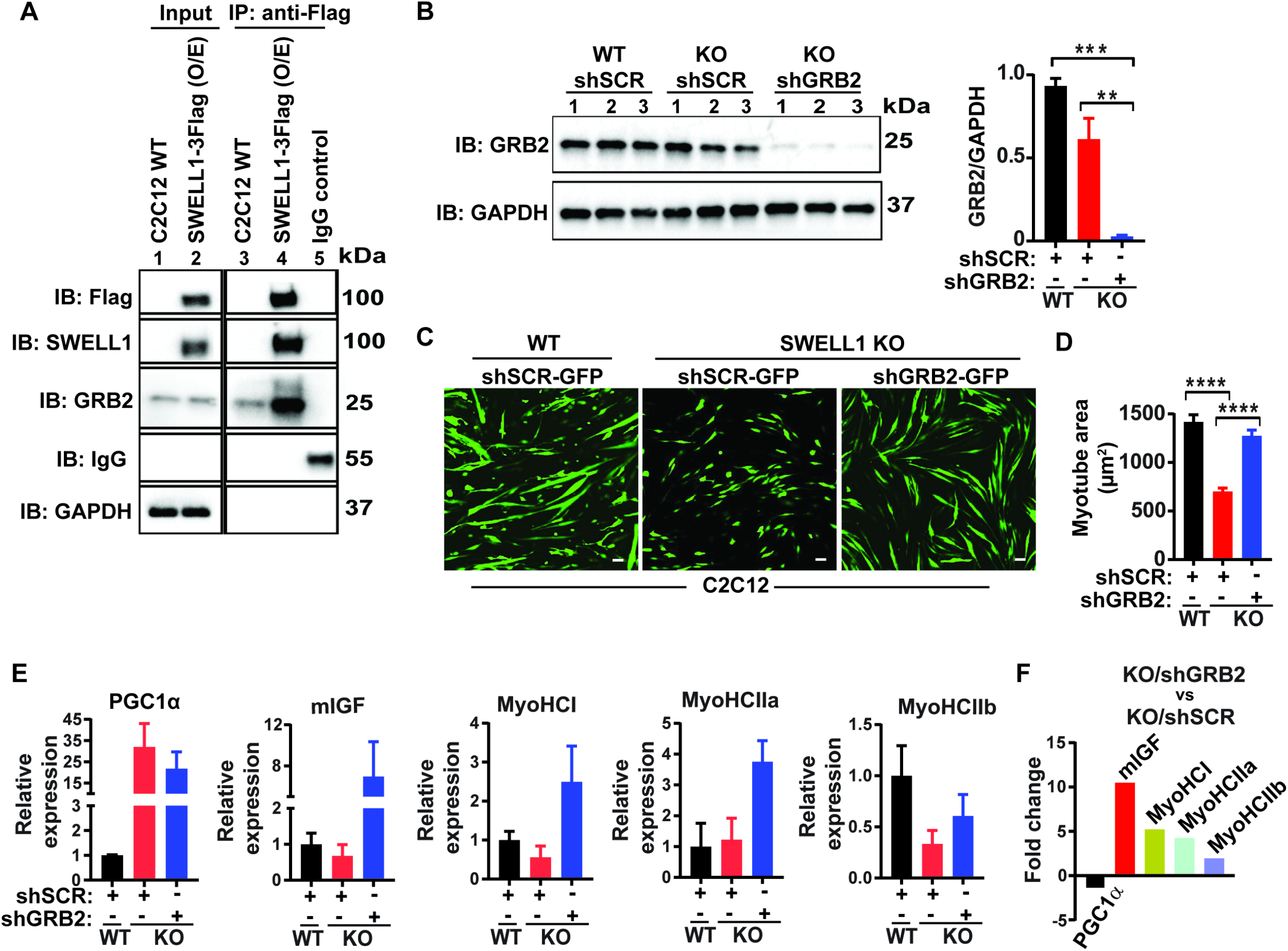
SWELL1 interacts with GRB2 in C2C12 myotubes and regulates myogenic differentiation. **A**, SWELL1-3xFlag over expressed in C2C12 cells followed by immunoprecipitation (IP) with Flag antibody. Western blot of Flag, SWELL1, GRB2 and GAPDH. IgG serves as a negative control. **B**, Western blot of GRB2 to validate GRB2 knock down efficiency in SWELL1 KO/GRB2 knock-down (Ad-shGRB2-GFP) compared to WT C2C12 (Ad-shSCR-GFP) and SWELL1 KO (Ad-shSCR-GFP). Densitometric quantification of GRB2 knock-down relative to GAPDH (right). **C**, Fluorescence image of WT C2C12/shSCR-GFP, SWELL1 KO/shSCR-GFP and SWELL1 KO/shGRB2-GFP myotubes. Scale bar: 100 μm. **D**, Quantification of mean myotube area of WT C2C12/shSCR-GFP (n=25), SWELL1 KO/shSCR-GFP (n=28) and SWELL1 KO/shGRB2-GFP (n=24). **E**, Relative mRNA expression of selected myogenic differentiation genes in SWELL1 KO/shSCR and SWELL1 KO/shGRB2 compared to WT C2C12/shSCR (n=3 each), and of SWELL1 KO/shGRB2 compared to SWELL1 KO/shSCR **(F)**, fold change of mRNA’s in KO shGRB2 relative to KO cells with preserved GRB2 expression. Statistical significance between the indicated group were calculated with one-way Anova, Tukey’s multiple comparisons test. Error bars represent mean ± s.e.m. *, P < 0.05, **, P < 0.01, ***, P < 0.001, ****, P < 0.0001. n = 3, independent experiments.

### Skeletal muscle targeted SWELL1 knock-out mice have reduced skeletal myocyte size, muscle endurance and ex vivo force generation

To examine the physiological consequences of SWELL1 ablation *in vivo*, we generated skeletal muscle specific SWELL1 KO mice using Cre-LoxP technology by crossing *Myf5-Cre* mice with *SWELL1^fl/fl^* mice (*Myf5 KO*; **Figure 6A**), and confirmed robust skeletal muscle SWELL1 depletion in *Myf5 KO* gastrocnemius muscle, 12.3-fold lower than WT controls (**Figure 6B**). Remarkably, in contrast to the severe impairments in skeletal myogenesis observed in both SWELL1 KO C2C12 and primary skeletal myotubes *in vitro* (**Figure 1,3,5**), *Myf5 KO* develop skeletal muscle mass comparable to WT littermates, based on Echo/MRI body composition (**Figure 6C**) and gross muscle weights (**Figure 6D**), and are born at normal mendelian ratios (**Supplementary file 3**). However, histological examination reveals a 27% reduction in skeletal myocyte cross-sectional area in *Myf5 KO* as compared to WT (**Figure 6E**), suggesting a requirement for *SWELL1* in skeletal muscle cell size regulation *in vivo*. This is potentially due to reductions in myotube fusion, as observed in C2C12 and primary skeletal muscle cells *in vitro* (**Figure 1**), but occurring to a lesser degree *in vivo*. These data indicate that the profound impairments in myogenesis observed *in vitro* may reflect a very early requirement for SWELL1 signaling in skeletal muscle development (prior to SWELL1 protein elimination by *Myf5-Cre* mediated *SWELL1* recombination), or other fundamental differences in myogenic differentiation processes *in vitro* versus *in vivo*.

**Figure 6:**
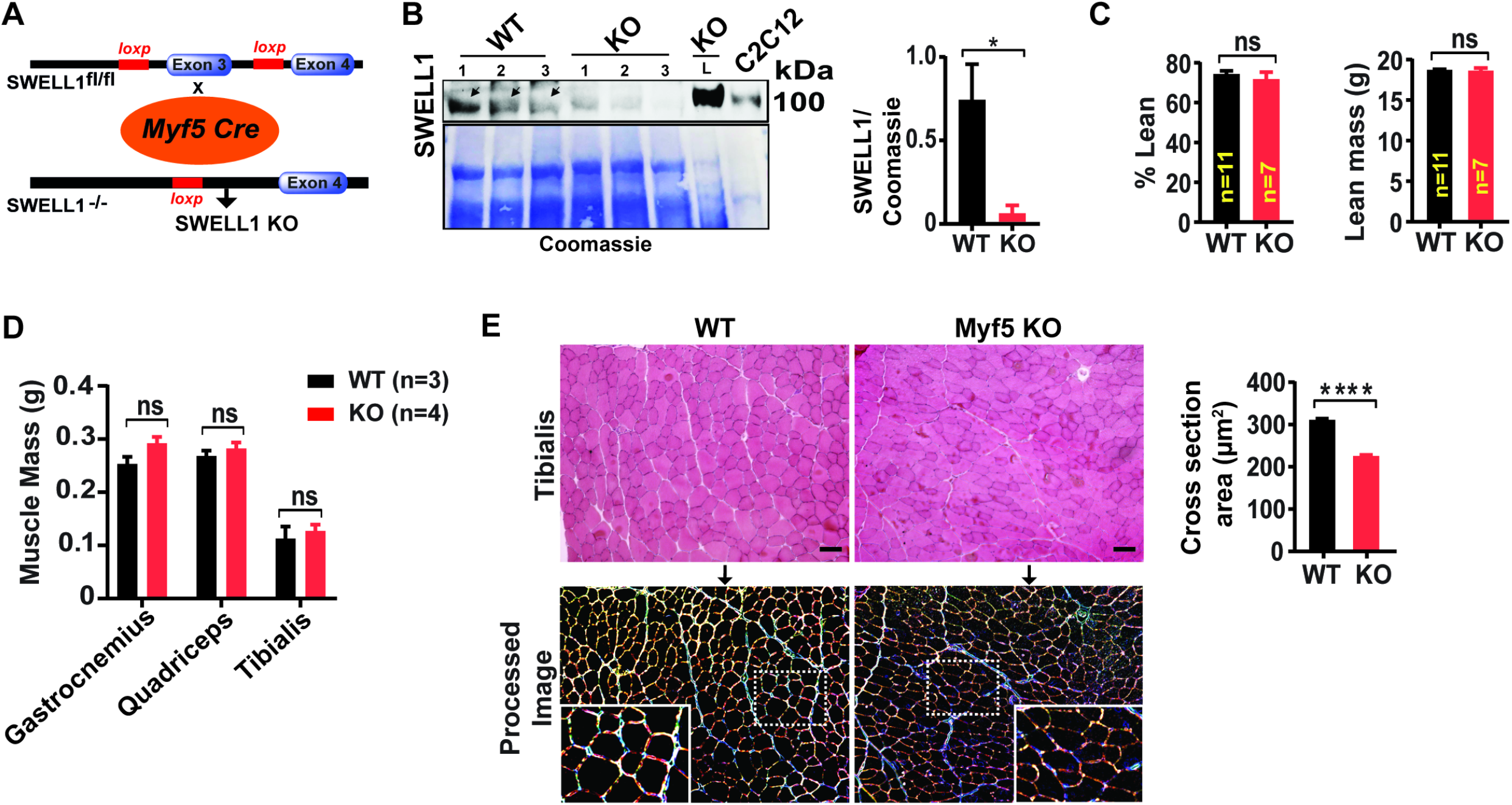
Skeletal muscle targeted SWELL1 KO mice develop smaller myofibers but normal muscle mass. **A**, Schematic representation of Cre-mediated recombination of loxP sites flanking Exon 3 using muscle-specific *Myf5-Cre* mice to generate skeletal muscle targeted SWELL1 KO mice. **B**, Western blot of gastrocnemius muscle protein isolated from of WT and *Myf5-Cre;SWELL1^fl/fl^* (*Myf5 KO*) mice. Liver sample from *Myf5 KO* and C2C12 cell lysates used as a positive control for SWELL1. Coomassie gel, below, serves as loading control for skeletal muscle protein. Densitometric quantification for SWELL1 deletion in skeletal muscle of *Myf5 KO* mice (n = 3) compared to WT (n = 3; *SWELL1^fl/fl^*) (right). **C**, NMR measurement of lean mass (%) and absolute fat mass of WT (n=11) and *Myf5 KO* (n=7) mice. **D**, Absolute muscle mass of muscle groups freshly isolated from WT (n=3) and *Myf5 KO* (n=4). **E**, Haematoxylin and eosin staining of tibialis muscle of WT and *Myf5 KO* mice fed on regular chow diet for 28 weeks (above). Scale bar: 100 μm. Below, ImageJ converted image highlights distinct surface boundaries of myotubes. Inset, enlarged image shows smaller fiber size in *Myf5 KO* muscle tissue. Quantification of average cross-sectional area of muscle fiber of WT (n=300) and *Myf5 KO* (n=300) mice from 10-12 different view field images (right). Statistical significance between the indicated values were calculated using a two-tailed Student’s t-test. Error bars represent mean ± s.e.m. *, P < 0.05, **, P < 0.01, ***, P < 0.001, ****, P < 0.0001.

Since insulin signaling is an important regulator of skeletal muscle oxidative capacity and endurance^47^, we next examined exercise tolerance on treadmill testing in *SWELL1^fl/fl^* (WT) compared to *Myf5 KO* mice. *Myf5 KO* mice exhibit a 14% reduced exercise capacity, compared to age and gender matched WT controls (**Figure 7A; Figure 7-Video 1**). Hang-times on inversion testing are also reduced 29% in *Myf5 KO* compared to controls, further supporting reduced skeletal muscle endurance upon skeletal muscle SWELL1 depletion *in vivo* (**Figure 7B**). To determine if these reductions in muscle function *in vivo* are due to muscle-specific functional impairments, we next performed *ex vivo* experiments in which we isolated the soleus muscle from mice and performed twitch/train testing. We observed that peak developed tetanic tension is 15% reduced in *Myf5 KO* soleus muscle compared to WT controls (**Figure 7C**), suggesting a skeletal muscle autonomous mechanism, with no difference in time to fatigability (TTF, **Figure 7D**) or time to 50% decay (**Figure 7E**).

**Figure 7:**
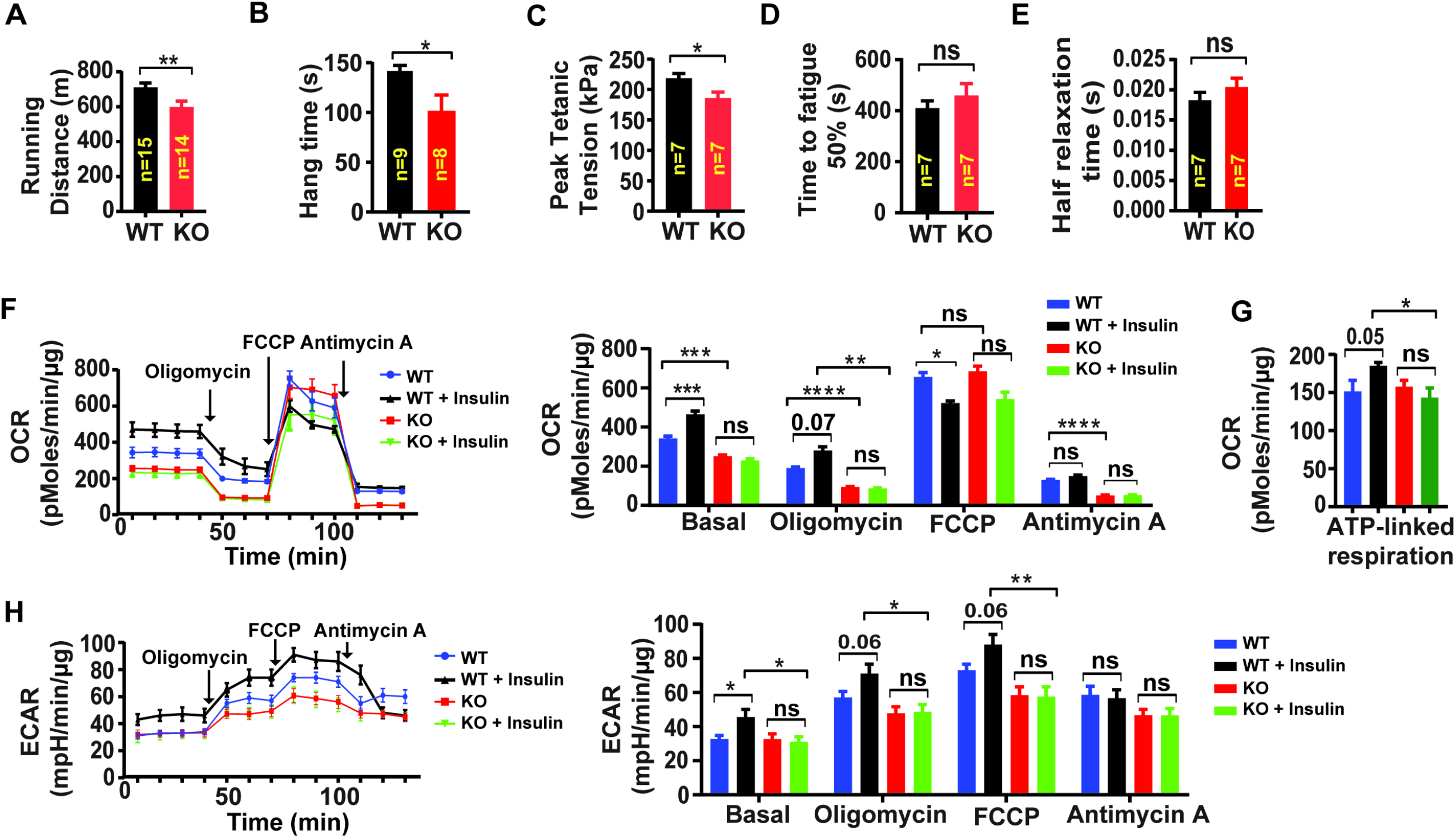
Skeletal muscle targeted SWELL1 deletion impairs muscle endurance, force generation and insulin-stimulated oxygen consumption. **A**, Exercise treadmill tolerance test for *Myf5 KO* mice (n = 14) compared to WT littermates (n=15). **B**, Hang times on inversion testing of *Myf5 KO* (n = 8) and WT (n= 9) mice. **C-E**, *Ex-vivo* isometric peak tetanic tension (**C**), time to fatigue (**D**) and half relaxation time (**E**) of isolated soleus muscle from *Myf5 KO* (n = 7) compared to WT (n = 7) mice. **F**, Oxygen Consumption Rate (OCR) in WT and SWELL1 KO primary myotubes +/- insulin stimulation (10 nM) (n=6 independent experiments) and quantification of basal OCR, OCR post Oligomycin, OCR post FCCP and OCR post Antimycin A. **G**, ATP-linked respiration obtained by subtracting the OCR after oligomycin from baseline cellular OCR. **H**, Extracellular acidification rate (ECAR) in WT and SWELL1 KO primary myotubes +/- insulin stimulation (10 nM) (n=6 independent experiments) and quantification of basal OCR, OCR post Oligomycin, OCR post FCCP and OCR post Antimycin A. Statistical significance between the indicated values were calculated using a two-tailed Student’s t-test. Error bars represent mean ± s.e.m. *, P < 0.05, **, P < 0.01, ***, P < 0.001, ****, P < 0.0001.

To determine whether these SWELL1 dependent differences in muscle endurance and force were due to impaired oxidative capacity, we next measured oxygen consumption rate (OCR) and extracellular acidification rate (ECAR) in WT and SWELL1 KO primary skeletal muscle cells, under basal and insulin-stimulated conditions (**Figure 7F**). Oxygen consumption of SWELL1 KO primary myotubes are 26% lower than WT and, in contrast to WT cells, are unresponsive to insulin-stimulation (**Figure 7F**), consistent with abrogation of insulin-AKT/ERK1/2 signaling upon skeletal muscle SWELL1 depletion. These relative changes persist in the presence of Complex V and III inhibitors, Oligomycin and Antimycin A (**Figure 7F&G**), suggesting that insulin-stimulated glycolytic pathways are primarily dysregulated upon SWELL1 depletion. In contrast, FCCP, which maximally uncouples mitochondria, abolishes differences in oxygen consumption between WT and SWELL1 KO primary muscle cells, suggesting that there might be no differences in functional mitochondrial content in SWELL1 KO muscle. To more directly measure glycolysis, we measured extracellular acidification rate (ECAR) in WT and SWELL1 KO primary myotubes. Insulin-stimulated ECAR increases are abolished in SWELL1 KO compared to WT cells, and these differences persist independent of electron transport chain modulators (**Figure 7H**). These data suggest that SWELL1 regulation of skeletal muscle cellular oxygen consumption occurs at the level of glucose metabolism - potentially via SWELL1-dependent insulin-PI3K-AKT-AS160-GLUT4 signaling, glucose uptake and utilization. These findings in primary skeletal muscle cells are supported by marked transcriptional suppression numerous glycolytic genes: Aldoa, Eno3, GAPDH, Pfkm, and Pgam2; and glucose and glycogen metabolism genes: Phka1, Phka2, Ppp1r3c and Gys1, upon SWELL1 ablation in C2C12 myotubes (**Figure 7-Figure supplement-1**).

### Skeletal muscle targeted SWELL1 ablation impairs systemic glucose metabolism and increases adiposity

Guided by evidence of impaired insulin-PI3K-AKT-AS160-GLUT4 signaling observed in SWELL1 KO C2C12 and primary myotubes we next examined systemic glucose homeostasis and insulin sensitivity in WT and *Myf5 KO* mice by measuring glucose and insulin tolerance. On a regular chow diet, there are no differences in either glucose tolerance or insulin tolerance (**Figure 8A**) between WT and *Myf5 KO* mice. However, over the course of 16-24 weeks on chow diet *Myf5 KO* mice develop 29% increased adiposity based on body composition measurements (**Figure 8B**) compared to WT, with no significant difference in lean mass (**Figure 6C**) or in total body mass (**Figure 8C**). When *Myf5 KO* mice are raised on a high-fat-diet (HFD) for 16 weeks there is no difference in adiposity observed (**Figure 8-Figure supplement-1**) compared to WT mice, but glucose tolerance is impaired (**Figure 8D**) and there is mild insulin resistance in HFD *Myf5 KO* mice as compared to WT (**Figure 8E**).

**Figure 8:**
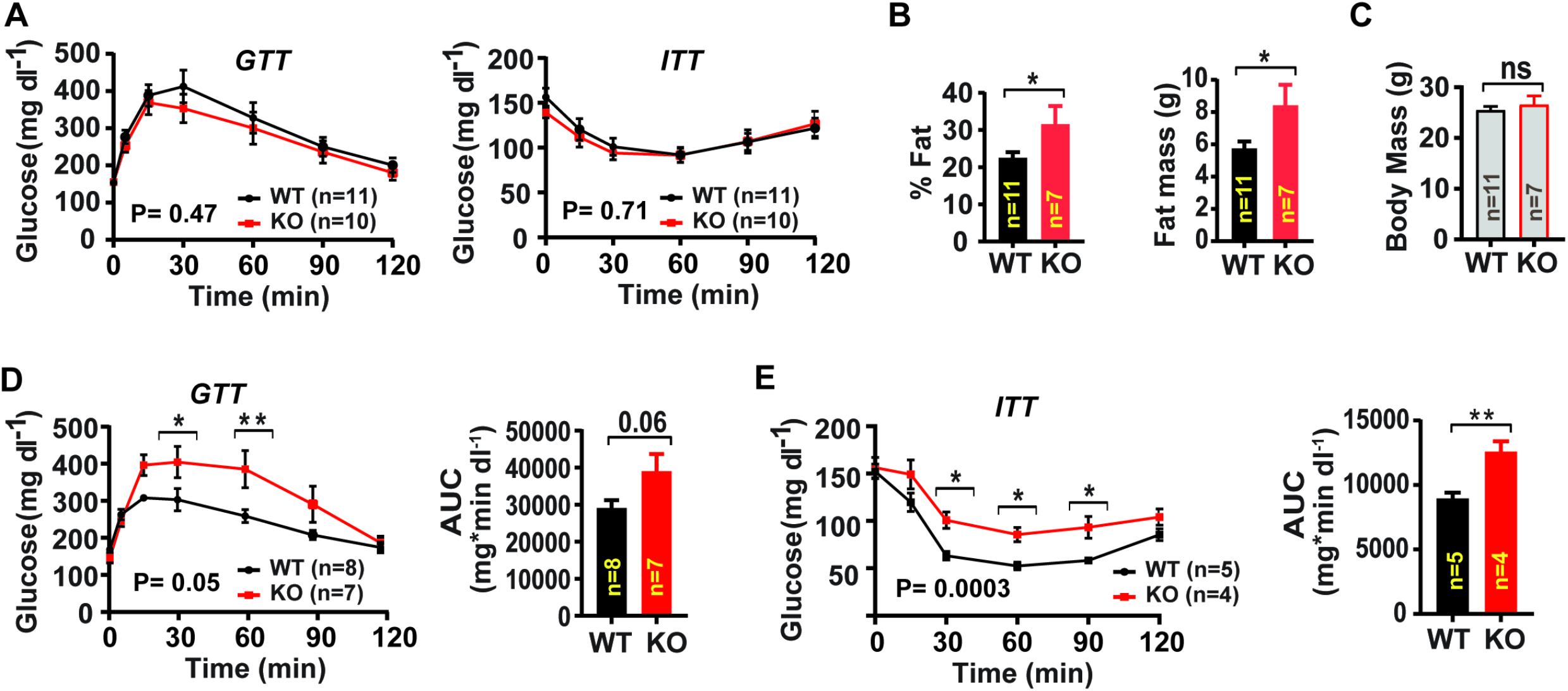
Skeletal muscle targeted SWELL1 ablation increases adiposity and induces glucose intolerance with overnutrition. **A**, Glucose and insulin tolerance tests of mice raised on chow diet of WT (n=11) and *Myf5 KO* (n=10) mice. **B**, NMR measurement of fat mass (%) and absolute fat mass of WT (n=11) and *Myf5 KO* (n=7) mice. **C**, Body mass of WT (n=11) and *Myf5 KO* (n=7) mice on regular chow diet. **D**, Glucose tolerance test of WT (n=8) and *Myf5 KO* (n=7) mice fed HFD for 16 weeks after 14-weeks of age. Corresponding area under the curve (AUC) for glucose tolerance for WT and *Myf5 KO* mice. **E**, Insulin tolerance tests of WT (n=5) and *Myf5 KO* (n=4) mice fed HFD for 18 weeks after 14-weeks of age. Corresponding area under the curve (AUC) for insulin tolerance for WT and *Myf5 KO* mice. Statistical significance test between the indicated group **B, C, D and E (AUC)** were calculated by using a two-tailed Student’s t-test. Error bars represent mean ± s.e.m. Two-way ANOVA was used for **A, D and E** (p-value in bottom corner of graph). Error bars represent mean ± s.e.m. *, P < 0.05, **, P < 0.01, ***, P < 0.001.

Since *Myf5* is also expressed in brown fat^48^, it is possible that these metabolic phenotypes arise from SWELL1-mediated effects in brown fat and consequent changes in systemic metabolism. To rule out this possibility, we repeated a subset of the above experiments in a skeletal muscle targeted KO mouse generated by crossing the *Myl1-Cre* and *SWELL1^fl/fl^* mice (*Myl1-Cre;SWELL1^fl/fl^) or Myl1 KO* (**Figure 8-Figure supplement-2A**), since *Myl1-Cre* is restricted to mature skeletal muscle (**Figure 8-Figure supplement-2B**), and excludes brown fat^49^. Similar to *Myf5 KO* mice, *Myl1 KO* mice fed a regular chow diet, have normal glucose tolerance (**Figure 8-Figure supplement-2C**), but exhibit 24% reduced exercise capacity on treadmill testing, as compared to WT (**Figure 8-Figure supplement-2D**). Also, *Myl1 KO* mice develop increased visceral adiposity over time on regular chow, based on 24% increased epididymal adipose mass normalized to body mass (**Figure 8-Figure supplement-2E**), with no differences in inguinal adipose tissue, muscle mass (**Figure 8-Figure supplement-2F**), or total body mass (**Figure 8-Figure supplement-2G**). These data suggest that impaired skeletal muscle glucose uptake in *Myl1 KO* and *Myf5 KO* mice are compensated for by increased adipose glucose uptake and *de novo* lipogenesis, which contribute to preserved glucose tolerance, at the expense of increased adiposity in skeletal muscle targeted SWELL1 KO mice raised on a regular chow diet. However, overnutrition-induced obesity, and the associated impairments in adipose and hepatic glucose disposal may uncover glucose intolerance and insulin resistance in skeletal muscle targeted SWELL1 KO mice.

## Discussion

Our data reveal that the SWELL1-LRRC8 channel complex regulates insulin/stretch-mediated AKT-AS160-GLUT4, MAP kinase and mTOR signaling in differentiated myoblast cultures, with consequent effects on myogenic differentiation, insulin-stimulated glucose metabolism and oxygen consumption. *In vivo*, skeletal muscle targeted SWELL1 KO mice have smaller skeletal muscle cells, impaired muscle endurance, and force generation, and are predisposed to adiposity, glucose intolerance and insulin resistance. Insulin/stretch-mediated PI3K-AKT, mTOR signaling are well known to be important regulators of myogenic differentiation ^50,51^, metabolism and muscle function ^2^ suggesting impaired SWELL1-AKT-mTOR signaling may underlie the defect in myogenic differentiation. Indeed, consistent with our previous findings and proposed model in adipocytes, in which SWELL1 mediates the interaction of GRB2 with IRS1 to regulate insulin-AKT signaling ^23,24^, SWELL1 also associates with GRB2 in skeletal myotubes, and GRB2 knock-down rescues impaired myogenic differentiation in SWELL1 KO muscle cells. Thus, our working model for SWELL1 mediated regulation of insuln-PI3K-AKT and downstream signaling in adipocytes ^24^ appears to be conserved in skeletal myotubes. The *in vitro* phenotype that we observe in CRISPR/cas9 mediated SWELL1 KO C2C12 myotubes and in SWELL1 KO primary myotubes is consistent with the observation of Chen et al (2019)^52^ that used siRNA mediated SWELL1 knock-down to demonstrate that the SWELL1-LRRC8 channel complex is required for myogenic differentiation. However, the ability of both GRB2 KD and SWELL1 O/E to rescue myogenic differentiation and augment insulin-AKT, MAP kinase and mTOR signaling in SWELL1 KO myotubes implicates non-canonical, non-conductive signaling mechanisms. Based on our work and also previous studies ^11,12^, SWELL1 O/E does not increase I_CI,SWELL_/VRAC to supranormal levels, although pAKT, pERK1/2 and mTOR levels are augmented by 2-fold to 3-fold above endogenous levels, upon 2-fold SWELL1 O/E in C2C12 myotubes. These data suggest that alternative/non-canonical signaling mechanisms underlie SWELL1-LRRC8 signaling, as opposed to canonical/conductive signaling mechanisms.

Another finding that warrants further study is the requirement for SWELL1 in stretch-induced AKT and MAP kinase signaling in C2C12 myotubes upon static stretching. Mechanical stretch is known to regulate myoblast proliferation and differentiation and myofiber hypertrophy via PI3K-AKT-MAPK signaling^2,42,43^. Also, there are numerous reports that VRAC, and presumably SWELL1-LRRC8, is mechano-responsive^25,26,13,37–41^. Therefore, it may not be surprising that SWELL1-LRRC8 complexes co-regulate both insulin and stretch-mediated PI3K-AKT, ERK1/2 signaling in skeletal myotubes - potentially integrating mechanical and hormonal stimuli to tune downstream signaling. Indeed, the concept of mechano-tuning insulin signaling has been proposed and demonstrated in other cell systems^53,54^ which implicate integrin signaling^54^ as the mechano-sensory mechanism. Curiously, it has been reported that VRAC can be activated in cardiac muscle cells by applying mechanical tension to β1-integrins ^25,26^, supporting the notion that, in striated muscle, integrin-SWELL1-LRRC8 may indeed participate in mechano-tuning insulin-AKT and downstream signaling. Further studies to more fully delineate the putative molecular mechanisms are necessary.

It is also notable that the profound myogenic differentiation block observed upon SWELL1 ablation in both C2C12 myotubes and primary myotubes *in vitro* is significantly milder *in vivo*, where only a 30% reduction in skeletal myocyte cross-sectional area is observed, with no change in total muscle mass, or lean content, in *Myf5 KO* mice. This discordance in phenotype may reflect fundamental differences in the biology of skeletal muscle differentiation *in vitro* versus the *in vivo milieu*. Alternatively, it may be that, although early, the time interval between *Myf5 Cre* expression early in myogenesis, and ultimate reductions in SWELL1 protein (potentially ~3 days) may extend beyond the critical period during which SWELL1 is required for myogenic differentiation. Directly testing this hypothesis would require examining mice expressing Cre-recombinase at the precursor stage, in skeletal muscle satellite cells, such as Pax3 or Pax7 promoters^55,56^ - these experiments are currently underway.

Although overall muscle development is grossly intact in both *Myl1 KO* and *Myf5 KO* mice, there is a consistent reduction in exercise capacity, muscle endurance and force generation, and a propensity for increased adiposity over time compared to age and gender matched controls. The observed impairments in exercise capacity in skeletal muscle SWELL1 KO mice are consistent with some level of insulin resistance, as in *db/db* mice^57^ and in humans^58^, and may be due to impaired skeletal muscle glycolysis and oxygen consumption in SWELL1 depleted skeletal muscle. Furthermore, the increased gonadal adiposity, with preserved glucose and insulin tolerance, observed in *Myl1 KO* and *Myf5 KO* mice phenocopy both skeletal muscle specific insulin receptor KO mice (MIRKO)^27^ and transgenic mice expressing a skeletal muscle dominant-negative insulin receptor mutant^28^, wherein skeletal muscle specific insulin resistance drives re-distribution of glucose from skeletal muscle to adipose tissue, to promote adiposity^29^. In the case of *Myf5 KO* mice, overnutrition and HFD feeding unmasks this underlying mild insulin resistance and glucose intolerance. Recent findings from skeletal muscle specific AKT1/AKT2 double KO mice indicate that these effects may not attributable to solely to muscle AKT signaling ^59^, but potentially involve other insulin sensitive signaling pathways.

In summary, we show that SWELL1-LRRC8 regulates myogenic differentiation and insulin-PI3K-AKT-AS160, ERK1/2, and mTOR signaling in myotubes via GRB2-mediated signaling. In vivo, SWELL1 is required for maintaining normal exercise capacity, muscle endurance, adiposity under basal conditions, and systemic glycemia in the setting of overnutrition. These finding contribute further to our understanding of SWELL1-LRRC8 channel complexes in the regulation of systemic metabolism.

## Author Contributions

Conceptualization, R.S.; methodology, A.K., L.X., C.M.T., A.H., S.K.G., R.M., K.S., J.M., C.G., R.S.; formal analysis, A.K., L.X., C.M.T, A.H, K.S., J.M.M., C.G., E.D.A., G.M., R.S.; investigation, A.K. L.X., A.H., S.K.G., R.M., K.S., C.M.T. R.S; resources, C.G., E.D.A., G.M., R.S.; writing (original draft), A.K., R.S., writing (review and editing), R.S., A.K., C.G., E.D.A., G.M.; visualization, A.K., C.M.T., L.X., S.K.G., A.H., R.M., C.G., R.S.; supervision, R.S., C.G., E.D.A., G.M.,; funding acquisition, C.G., G.M. E.D.A., R.S.

## Acknowledgements

RNA-Seq data presented herein were obtained at the Genomics Division of the Iowa Institute of Human Genetics. This work was supported by grants NIH/NHLBI R01 HL125436 (C.E.G), NIH RO1HL127764, RO1HL112413 (E.D.A), NIH NIDDK 1R01DK106009 (R.S.), the Roy J. Carver Trust (R.S.) and Musculoskeletal Research Center Pilot Grant 1P30AR074992-01 (R.S.).

## Supplementary Figure Legends

**Figure 7-Figure supplement-1:**
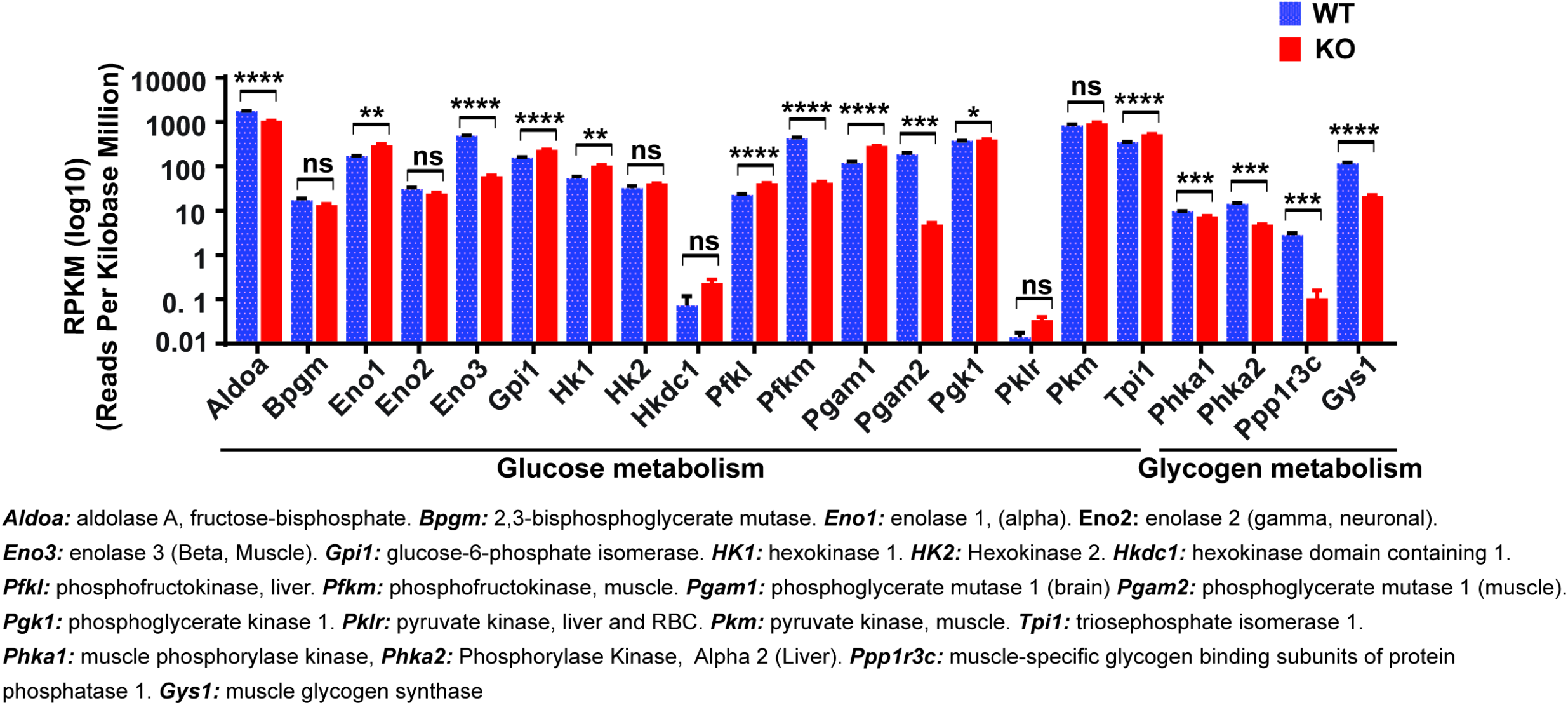
SWELL1 ablation in C2C12 myotubes downregulates glucose and glycogen associated genes. Differentially expressed glucose and glycogen metabolism associated gene after RNA-seq analysis of C2C12 WT and SWELL1 KO myotube (n= 3, each). Statistical significance between the indicated values were calculated using a two-tailed Student’s t-test. Error bars represent mean ± s.e.m. *, P < 0.05, **, P < 0.01, ***, P < 0.001, ****, P < 0.0001.

**Figure 8-Figure supplement-1:**
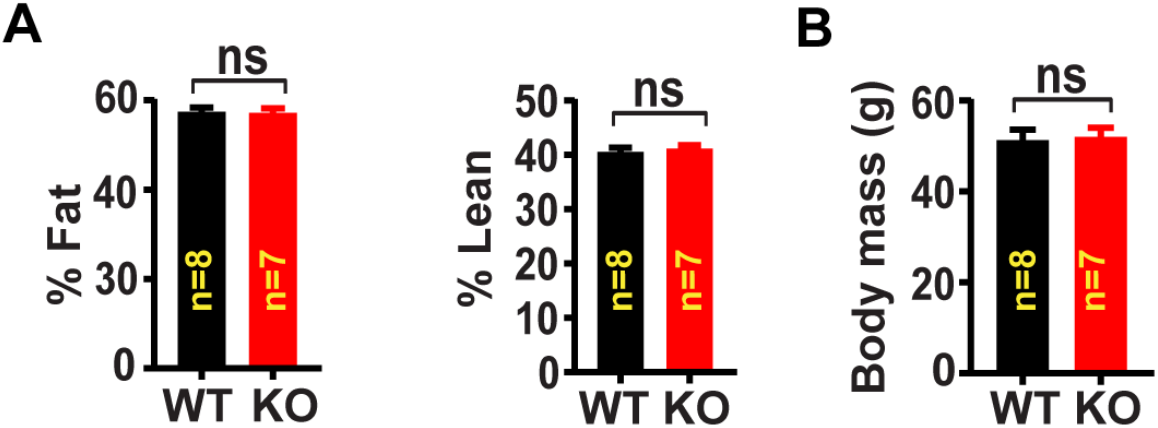
Body composition of skeletal muscle specific SWELL1 KO mice raised on a high-fat-diet (HFD). **A**, NMR measurement of fat mass (%) and lean mass (%) of WT (n=8) and *Myf5 KO* (n=7) mice raised on HFD (16 weeks) after 14-weeks of age. **B**, Body mass of WT (n=8) and *Myf5 KO* (n=7) mice.

**Figure 8-Figure supplement-2:**
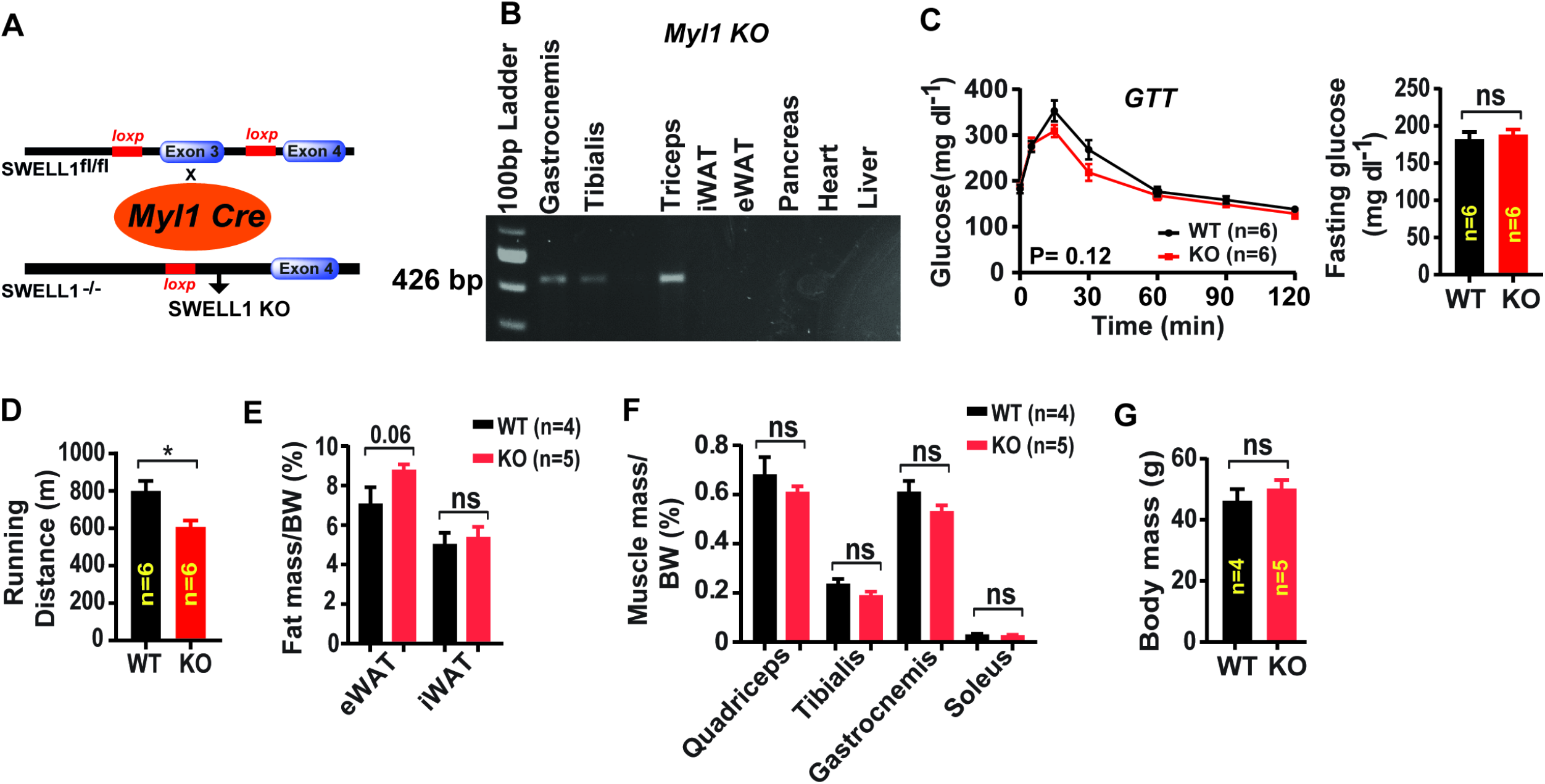
Skeletal muscle targeted SWELL1 ablation impairs muscle endurance and induces adiposity. **A**, Schematic representation of Cre-mediated recombination of loxP sites flanking Exon 3 using muscle-specific *Myl1-Cre* mice to generate skeletal muscle targeted SWELL1 KO mice (*Myl1-Cre;SWELL1^fl/fl^; Myl1 KO*) **B**, PCR band of SWELL1 recombination in *Myl1 KO* mice from isolated tissues. **C**, Glucose tolerance test of WT (n=6) and *Myl1KO* (n=6) mice raised on chow food diet for 14 weeks. Fasting glucose level for WT and *Myl1KO* mice (right). **D**, Exercise treadmill tolerance test for *Myl1KO* (n=6) compared to WT (n=6) littermates. **E**, Epididymal (eWAT) and inguinal (iWAT) fat mass normalized to body mass (BM) isolated from *Myl1 KO* (n=5) and WT (n=4) mice. **F**, Skeletal muscle mass normalized to body mass (BM) isolated from *Myl1 KO* (n=5) and WT (n=4) mice. **G**, Body mass of *Myl1 KO* (n=5) and WT (n=4) mice raised on regular chow diet. Statistical significance between the indicated values were calculated using a two-tailed Student’s t-test. Error bars represent mean ± s.e.m. *, P < 0.05.

## Materials and methods

### Animals

The Institutional Animal Care and Use Committee of the Washington University in St. Louis and the University of Iowa approved all experimental procedures involving animals. All the mice were housed in temperature, humidity, and light-controlled room and allowed free access to water and food. Both male and female *SWELL1*^*fl/f*l^ (WT), *Myl1Cre;SWELL1^fl/fl^ (Myl1 KO), Myf5Cre;SWELL1^fl/fl^* (skeletal muscle targeted SWELL1 KO), were generated and used in these studies. *Myl1Cre* (JAX# 24713) and *Myf5Cre* (*JAX#* 007893) mice were purchased from Jackson labs. For high-fat diet (HFD) studies, we used Research Diets Inc. (Cat # D12492) (60 kcal% fat) regimen starting at 14 weeks of age.

### Generation of CRISPR/Cas9-mediated *SWELL1* floxed (*SWELL1^fl/fl^*) mice

*SWELL1^fl/fl^* mice were generated as previously described^1^. Briefly, *SWELL1* intronic sequences were obtained from Ensembl Transcript ID ENSMUST00000139454. All CRISPR/Cas9 sites were identified using ZiFit Targeter Version 4.2 (Http://zifit.partners.org/ZiFiT/). Pairs of oligonucleotides corresponding to the chosen CRISPR-Cas9 target sites were designed, synthesized, annealed, and cloned into the pX330-U6-Chimeric_BB-CBh-hSpCas9 construct (Addgene plasmid # 42230), following the protocol detailed in Cong et al. (2013)^2^. CRISPR-Cas9 reagents and ssODNs were injected into the pronuclei of F1 mixed C57/129 mouse strain embryos at an injection solution concentration of 5 ng/μl and 75-100 ng/μl, respectively. Correctly targeted mice were screened by PCR across the predicted loxP insertion sites on either side of Exon 3. These mice were then backcrossed >8 generations into a C57BL/6 background.

### Antibodies

Rabbit polyclonal anti-SWELL1 antibody was generated against the epitope QRTKSRIEQGIVDRSE (Pacific Antibodies). All other primary antibodies were purchased from Cells Signaling: anti-β-actin (#8457s), p-AKT1 (#9018s), Akt1 (#2938s), pAKT2 (#8599s), Akt2 (#3063s), p-AS160 (#4288s), AS160 (#2670s), AMPKα (#5831s), pAMPKα (#2535s), FoxO1(#2880s) and pFoxO1(#9464s), p70 S6 Kinase (#9202s), p-p70 S6 Kinase (#9205s), pS6 Ribosomal (#5364s), GAPDH (#5174s), pErk1/2 (#9101s), Total Erk1/2 (#9102s). Purified mouse anti-Grb2 was purchased from BD (610111s). Purified anti-flag mouse antibody was purchased from sigma. Rabbit IgG Santa Cruz (sc-2027). All primary antibodies were used at 1:1000 dilution, except for anti-flag at 1:2000 dilution. All secondary antibody (anti-rabbit-HRP and anti-mouse-HRP) were used at 1:10000 dilution.

### Adenovirus

Adenovirus type 5 with Ad5-CMV-mCherry (1 X 10^10^ PFU/ml), Ad5-CMV-Cre-mCherry (3 X 10^10^ PFU/ml) were obtained from the University of Iowa viral vector core facility. Ad5-CAG-LoxP-stop-LoxP-3XFlag-SWELL1 (1X 10^10^ PFU/ml) were obtained from Vector Biolabs. Ad5-U6-shGRB2-GFP (1 X 10^9^ PFU/ml) and Ad5-U6-shSCR-GFP (1 X 10^10^ PFU/ml) were obtained from Vector Biolabs.

### Electrophysiology

All recordings were performed in the whole-cell configuration at room temperature, as previously described^1,3^. Briefly, currents were measured with either an Axopatch 200B amplifier or a MultiClamp 700B amplifier (Molecular Devices) paired to a Digidata 1550 digitizer, using pClamp 10.4 software. The intracellular solution contained (in mM): 120 L-aspartic acid, 20 CsCl, 1 MgCl_2_, 5 EGTA, 10 HEPES, 5 MgATP, 120 CsOH, 0.1 GTP, pH 7.2 with CsOH. The extracellular solution for hypotonic stimulation contained (in mM): 90 NaCl, 2 CsCl, 1 MgCl_2_, 1 CaCl2, 10 HEPES, 5 glucose, 5 mannitol, pH 7.4 with NaOH (210 mOsm/kg). The isotonic extracellular solution contained the same composition as above except for mannitol concentration of 105 (300 mOsm/kg). The osmolarity was checked by a vapor pressure osmometer 5500 (Wescor). Currents were filtered at 10 kHz and sampled at 100 μs interval. The patch pipettes were pulled from borosilicate glass capillary tubes (WPI) using a P-87 micropipette puller (Sutter Instruments). The pipette resistance was ~4-6 MΩ when the patch pipette was filled with intracellular solution. The holding potential was 0 mV. Voltage ramps from −100 to +100 mV (at 0.4 mV/ms) were applied every 4 s.

### Primary muscle satellite cell isolation

Satellite cell isolation and differentiation were performed as described previously with minor modifications^4^. Briefly, gastrocnemius and quadriceps muscles were excised from *SWELL1^flfl^* mice (8-10 weeks old) and washed twice with 1XPBS supplemented with 1% penicillin-streptomycin and fungizone (300 ul/100ml). Muscle tissue was incubated in DMEM-F12 media supplemented with collagenase II (2 mg/ml), 1% penicillin-streptomycin and fungizone (300 ul/100ml) and incubated at shaker for 90 minutes at 37°C. Tissue was washed with 1X PBS and incubated again with DMEM-F12 media supplemented with collagenase II (1 mg/ml), dispase (0.5 mg/ml), 1% penicillin-streptomycin and fungizone (300ul/100ml) in a shaker for 30 minutes at 37°C. Subsequently, the tissue was minced and passed through a cell strainer (70 μm), and after centrifugation; satellite cells were plated on BD Matrigel-coated dishes. Cells were stimulated to differentiate into myoblasts in DMEM-F12, 20% fetal bovine serum (FBS), 40 ng/ml basic fibroblast growth factor (bfgf, R&D Systems, 233-FB/CF), 1X non-essential amino acids, 0.14 mM β-mercaptoethanol, 1X penicillin/streptomycin, and Fungizone. Myoblasts were maintained with 10 ng/ml bfgf and then differentiated in DMEM-F12, 2% FBS, 1X insulin–transferrin–selenium, when 80% confluency was reached.

### Cell culture

WT C2C12 and *SWELL1* KO C2C12 cell line were cultured at 37°C, 5% CO2 Dulbecco’s modified Eagle’s medium (DMEM; GIBCO) supplemented with 10% fetal bovine serum (FBS; Atlanta Bio selected) and antibiotics 1% penicillin-streptomycin (Gibco, USA). Cells were grown to 80% confluency and then transferred to differentiation media DMEM supplemented with antibiotics and 2% horse serum (HS; GIBCO) to induce differentiation. The differentiation media was changed every two days. Cells were allowed to differentiate into myotubes for up to 6 days. Subsequently, myotube images were taken for quantification of myotube surface area and fusion index.

### Myotube morphology, surface area and fusion index quantification

After differentiation (Day 7), cells were imaged with Olympus IX73 microscope (10X objective, Olympus, Japan). For each experimental condition, 5-6 bright field images were captured randomly from 6 well plate. Myotube surface area was quantified manually with ImageJ software. The morphometric quantification was carried out by an independent observer who was blinded to the experimental conditions. For fusion index, differentiated myotube growing on coverslip were washed with 1X PBS and fixed with 2% PFA. After washing with 1XPBS 3 times, cells were permeabilized with 0.1% TritonX100 for 5 minutes at room temperature and subsequently blocking was done with 5% goat serum for 30 minutes. Cells were stained with DAPI (1uM) for 15 minutes and after washing with 1X PBS, coverslip were mounted on slides with ProLong Diamond anti-fading agent. Cells were imaged with Olympus IX73 microscope (10X objective, Olympus, Japan) with bright field and DAPI filter. Fusion index (number of nuclei incorporated within the myotube/ total number of nuclei present in that view field) were analyzed by ImageJ.

### RNA sequencing

RNA quality was assessed by Agilent BioAnalyzer 2100 by the University of Iowa Institute of Human Genetics, Genomics Division. RNA integrity numbers greater than 8 were accepted for RNAseq library preparation. RNA libraries of 150 bp PolyA-enriched RNA were generated, and sequencing was performed on a HiSeq 4000 genome sequencing platform (Illumina). Sequencing results were uploaded and analyzed with BaseSpace (Illumina). Sequences were trimmed to 125 bp using FASTQ Toolkit (Version 2.2.0) and aligned to Mus musculus mmp10 genome using RNA-Seq Alignment (Version 1.1.0). Transcripts were assembled and differential gene expression was determined using Cufflinks Assembly and DE (Version 2.1.0). Ingenuity Pathway Analysis (QIAGEN) was used to analyze significantly regulated genes which were filtered using cutoffs of >1.5 fragments per kilobase per million reads, >1.5 fold changes in gene expression, and a false discovery rate of <0.05. Heatmaps were generated to visualize significantly regulated genes.

### Myotube signaling studies

For insulin stimulation, differentiated C2C12 myotubes were incubated in serum free media for 6 h and stimulated with 0 and 10 nM insulin for 15 min; while differentiated primary myotubes were incubated in serum free media for 4 h and stimulated with 0 and 10 nM insulin for 2 h. To examine intracellular signaling upon SWELL1 overexpression (SWELL1 O/E), we overexpressed SWELL1-3xFlag by transducing C2C12 myotubes with Ad5-CAG-LoxP-stop-LoxP-SWELL1-3xFlag (MOI 50-60) and Ad5-CMV-Cre-mCherry (MOI 50-60) and polybrene (4 μg/ml) in DMEM (2% FBS and 1% penicillin-streptomycin) for 36 h. Ad5-CMV-Cre-mCherry alone with polybrene (4 μg/ml) (MOI 50-60) was transduced in WT C2C12 or SWELL1 KO C2C12 as controls. Viral transduction efficiency (60-70%) was confirmed by mCherry fluorescence. Cells were allowed to differentiate further in differentiation media up to 6 days. Myotube images were taken before collecting lysates for further signaling studies. GRB2 knock-down was achieved by transducing myotubes with Ad5-U6-shSCR-GFP (Control, MOI 50-60) or Ad5-U6-shSWELL1-GFP (GRB2 KD, MOI 50-60) in DMEM (2% FBS and 1% penicillin-streptomycin) supplemented with polybrene (4 μg/ml) for 24 hour. Cells were allowed to differentiate further in differentiation media up to 6 days. Differentiated myotube images were taken for myotube surface area quantification before collecting the cells for RNA isolation.

### Stretch stimulation

C2C12 myotubes were plated in each well of a 6 well BioFlex culture plate. Cells were allowed to differentiate up to 6 days in differentiation media, and then placed into a Flexcell Jr. Tension System (FX-6000T) and incubated at 37°C with 5% CO2. C2C12 myotubes on flexible membrane were subjected to either no tension or to static stretch of 5% for 15 minutes. Cells were lysed and protein isolated for subsequent Western blots.

### Western blot

Cells were washed with ice cold 1X PBS and lysed in ice-cold lysis buffer (150 mM NaCl, 20 mM HEPES, 1% NP-40, 5mM EDTA, pH 7.5) with added proteinase/phosphatase inhibitor (Roche). The cell lysate was further sonicated (20% pulse frequency for 20 sec) and centrifuged at 14000 rpm for 20 min at 4°C. The supernatant was collected and estimated for protein concentration using DC protein assay kit (Bio-Rad). For immunoblotting, an appropriate volume of 4 x Laemmli (Bio-rad) sample loading buffer was added to the sample (10-20 μg of protein), then heated at 90°C for 5 min before loading onto 4-20% gel (Bio-Rad). Proteins were separated using running buffer (Bio-Rad) for 2 h at 110 V. Proteins were transferred to PVDF membrane (Bio-Rad) and membrane blocked in 5% (w/v) BSA or 5 % (w/v) milk in TBST buffer (0.2 M Tris, 1.37 M NaCl, 0.2% Tween-20, pH 7.4) at room temperature for 1 hour. Blots were incubated with primary antibodies at 4 °C overnight, followed by secondary antibody (Bio-Rad, Goat-anti-mouse #170-5047, Goat-anti-rabbit #170-6515, all used at 1:10000) at room temperature for one hour. Membranes were washed 3 times and imaged by chemiluminescence (Pierce) by using a Chemidoc imaging system (BioRad). The images were further analyzed for band intensities using ImageJ software. β- Actin or GAPDH levels were quantified for equal protein loading.

### Immunoprecipitation

C2C12 myotubes were plated on 10 cm dishes in complete media and grown to 80% confluency. For SWELL1-3xFlag overexpression, Ad5-CAG-LoxP-stop-LoxP-3XFlag-SWELL1 (MOI 50-60) and Ad5-CMV-Cre-mCherry (MOI 50-60) along with polybrene (4 ug/ml) were added to cells in DMEM media (2% FBS and 1% penicillin-streptomycin) allowed to grow for 36 hours. Cells were then switched to differentiation media for up to 6 days. After that myotubes were harvested in ice-cold lysis buffer (150 mM NaCL, 20 mM HEPES, 1% NP-40, 5mM EDTA, pH 7.5) with added protease/phosphatase inhibitor (Roche) and kept on ice with gentle agitation for 15 minutes to allow complete lysis. Lysated were incubated with anti-Flag antibody (Sigma #F3165) or control rabbit IgG (Santa Cruz sc-2027) rotating end over end overnight at 4°C. Protein G sepharose beads (GE) were added for 4 h and then samples were centrifuged at 10,000g for 3 minutes and washed three times with RIPA buffer and re-suspended in laemmli buffer (Bio-Rad), boiled for 5 minutes, separated by SDS-PAGE gel followed by the western blot protocol.

### RNA isolation and quantitative RT-PCR

Differentiated cells were solubilized in TRIzol and the total RNA was isolated using PureLink RNA kit (Life Technologies) and column DNase digestion kit (Life Technologies). The cDNA synthesis, qRT-PCR reaction and quantification were carried out as described previously^1^. All experiment was performed in triplicate and GAPDH were used as internal standard to normalize the data. All primers used for qRT-PCR are listed in **Supplementary file 4**.

### Muscle tissue homogenization

Mice were euthanized and gastrocnemius muscle excised and washed with 1X PBS. Muscles tissue were minced with surgical blade and kept in 8 volume of ice cold homogenization buffer (20 mM Tris, 137 mM NaCl, 2.7 mM KCl, 1 mM MgCl_2_, 1 % Triton X-100, 10 % (w/v) glycerol, 1 mM EDTA, 1 mM dithiothreitol, pH 7.8) supplemented with protease/phosphatase inhibitor (Roche). Tissues were homogenized on ice with a Dounce homogenizer (40–50 passes) and incubated for overnight at 4 °C with continuous rotation. Tissue lysate was further sonicated in 20 sec cycle intervals for 2-3 times and centrifuged at 14000 rpm for 20 min at 4°C. The supernatant was collected for protein concentration estimation using DC protein assay kit (Bio-Rad). Due to the high content of contractile protein in this preparation, coomassie gel staining was performed to demonstrate equal protein loading, and for quantification normalization of Western blots.

### Tissue histology

Mice were anesthetized with isoflurane followed by cervical dislocation. Tibialis anterior (TA) muscle was carefully excised and gently immersed into the tissue-tek O.C.T medium placed on wooden cork. Orientation of the tissue maintained while embedding in the medium. Subsequently, wooden cork with tissue gently immersed into the liquid N2 pre-chilled isopentane bath for 10-14 sec and store at −80°C. Tissue sectioning (10 μm) were done with Leica cryostat and all sections collected on positively charged microscope slide for H&E staining as described earlier^5^. Briefly, TA sectioned slides were stained for 2 minutes in hematoxylin, 1 minute in eosin and then dehydrated with ethanol and xylenes. Subsequently, slides were mounted with coverslip and image were taken with EVOS cell imaging microscope (10X objective). For quantification of fiber cross-sectional area, images were processed using ImageJ software to enhance contrast and smooth/sharpen cell boundaries and clearly demarcate muscle fiber cross sectional area. All measurement was performed with an independent observer who was blinded to the identity of the slides.

### Exercise tolerance test and inversion testing

Mouse treadmill exercise protocols were adapted from Dougherty et al. (2016)^6^. Briefly, mice were first acclimated with the motorized treadmill (Columbus Instruments Exer3/6 Treadmill (Columbus, OH) for 3 days by running 10-15 minutes (with 3 minutes interval) for 3 consecutive days at 7 m/min, with the electric shocking grid (frequency 1 Hz) installed in each lane. During the treadmill testing, mice ran with a gradual increase in speed (5.5 m/minute to 22 m/minute) and inclination (0°-15°) at time intervals of 3 minutes each. The total running distance for each mouse was recorded at the end of the experiment. The predefined criteria for removing the mouse from the treadmill and recording the distance travelled was: continuous shock for 5 sec or receiving 5-6 shocks within a time interval of 15 seconds. These mice were promptly removed from the treadmill and total duration and distance were recorded for further analysis.

Mouse inversion test was performed using a wire-grid screen apparatus elevated to 50 cm. Mice were stabilized on the screen inclined at 60°, with the mouse head facing towards the base of the screen. The screen was slowly pivoted to 0° (horizontal), such that the mouse was fully inverted and hanging upside down from the screen. Soft bedding was placed underneath the screen to protect mouse from any injury, were they to fall. The inversion test for each mouse was repeated 2 times with an interval of 45 minutes (resting period). The hang time for each mouse was repeated 3 times with an interval of 5-minute. The maximum hanging time limit for each mouse was set for 3 minutes.

### Isolated muscle contractile assessment

Soleus muscle was carefully dissected and transferred to a specialized muscle stimulation system (1500A, Aurora Scientific, Aurora, ON, Canada) where physiology tests were run in a blinded fashion. Muscle was immersed in a Ringer solution (in mM) (NaCl 137, KCI 5, CaCl_2_ 2, NaH_2_PO_4_ 1, NaHCO_3_ 24, MgSO_4_ 1, glucose 11 and curare 0.015) maintained at 37°C. The distal tendon was secured with silk suture to the arm of a dual mode ergometer (300C-LR, Aurora Scientific, Aurora, ON, Canada) and the proximal tendon secured to a stationary post. Muscles were stimulated with an electrical stimulator (701C, Aurora Scientific, Aurora, ON, Canada) using parallel platinum plate electrodes extending along the muscle. Muscle slack length was set by increasing muscle length until passive force was detectable above the noise of the transducer and fiber length was measured through a micrometer reticule in the eyepiece of a dissecting microscope. Optimal muscle length was then determined by incrementally increasing the length of the muscle by 10% of slack fiber length until the isometric tetanic force plateaued. At this optimum length, force was recorded during a twitch contraction and isometric tetanic contraction (300 ms train of 0.3 ms pulses at 225 Hz). The muscle was then fatigued with a bout of repeated tetanic contractions every 10 seconds until force dropped below 50% of peak. At this point, the muscle was cut from the sutures and weighed. This weight, along with peak fiber length and muscle density (1.056 g/cm3), was used to calculate the physiological cross-sectional area (PCSA) and convert to specific force (tension). The experimental data were analyzed and quantified using Matlab (Mathworks), and presented as peak tetanic tension (Tetanic Tension) – peak of the force recording during the tetanic contraction, normalized to PCSA; Time to fatigue (TTF) – time for the tetanic tension to fall below 50% of the peak value during the fatigue test; Half relaxation time (HRT) – half the time between force peak and return to baseline during the twitch contraction.

### XF-24 Seahorse assay

Cellular respiration was quantified in primary myotubes using the XF24 extracellular flux (XF) bioanalyzer (Agilent Technologies/Seahorse Bioscience, North Billerica, MA, USA). Primary skeletal muscle cells isolated from *SWELL1^flfl^* mice were plated on BD Matrigel-coated plate at a density of 20 × 10^3^ per well. After 24 hours, cells were incubated in Ad5-CMV-mCherry or Ad5-CMV-Cre-mCherry (MOI 90-100) in DMEM-F12 media (2% FBS and 1% penicillin-streptomycin) for 24 hours. Cells were then switched to differentiation media for another 3 days. For insulin-stimulation, cells were incubated in serum free media for 4 h and stimulated with 0 and 10 nM insulin for 2 h. Subsequently, medium was changed to XF-DMEM, and kept in a non-CO_2_ incubator for 60 minutes. The basal oxygen consumption rate (OCR) was measured in XF-DMEM. Subsequently, oxygen consumption was measured after addition of each of the following compounds: oligomycin (1 μg/ml) (ATP-Linked OCR), carbonyl cyanide 4-(trifluoromethoxy) phenylhydrazone (FCCP; 1 μM) (Maximal Capacity OCR) and antimycin A (10 μM; Spare Capacity OCR)^7^. For the glycolysis stress test, prior to experimentation, cells were switched to glucose-free XF-DMEM and kept in a non-CO_2_ incubator for 60 min. Extracellular acidification rate (ECAR) was determined in XF-DMEM followed by these additional conditions: glucose (10 mM), oligomycin (1 μM), and 2-DG (100 mM). Data for Seahorse experiments (normalized to protein) reflect the results of one Seahorse run/condition with 6 replicates.

### Metabolic phenotyping

Mouse body composition (fat and lean mass) was measured by nuclear magnetic resonance (NMR); Echo-MRI 3-in-1 analyzer, EchoMRI, LLC). For glucose tolerance test (GTT), mice were fasted for 6 hours and intraperitoneal injection of glucose (1g/kg body weight for lean mice and 0.75g/kg of body weight for HFD mice) administered. Glucose level was monitored from tail-tip blood using a glucometer (Bayer Healthcare LLC) at the indicated times. For insulin tolerance test (ITT), mice were fasted for 4 hours and after an intra-peritoneal injection of insulin (HumulinR, 1U/kg for lean mice and 1.25U/kg for HFD mice) glucose level was measured by glucometer at the indicated times.

### Statistics

Data are represented as mean ± s.e.m. Two-tail paired or unpaired Student’s t-tests were used for comparison between two groups. For three or more groups, data were analyzed by one-way analysis of variance and Tukey’s *post hoc* test. For GTTs and ITTs, 2-way analysis of variance (Anova) was used. A p-value < 0.05 was considered statistically significant. *, ** and *** represents a p-value less than 0.05, 0.01 and 0.001 respectively.

## Notes

### Competing Interest Statement

The authors have declared no competing interest.

## References

1. Fitts, R.H., Riley, D.R. & Widrick, J.J. Functional and structural adaptations of skeletal muscle to microgravity. J Exp Biol 204, 3201–3208 (2001).

2. Schiaffino, S., Dyar, K.A., Ciciliot, S., Blaauw, B. & Sandri, M. Mechanisms regulating skeletal muscle growth and atrophy. FEBS J 280, 4294–4314 (2013).

3. Ben-Sahra, I. & Manning, B.D. mTORC1 signaling and the metabolic control of cell growth. Curr Opin Cell Biol 45, 72–82 (2017).

4. Hornberger, T.A., Sukhija, K.B. & Chien, S. Regulation of mTOR by mechanically induced signaling events in skeletal muscle. Cell cycle 5, 1391–1396 (2006).

5. Yoon, M.S. mTOR as a Key Regulator in Maintaining Skeletal Muscle Mass. Frontiers in physiology 8, 788 (2017).

6. Carson, J.A. & Wei, L. Integrin signaling’s potential for mediating gene expression in hypertrophying skeletal muscle. J Appl Physiol (1985) 88, 337–343 (2000).

7. Schlaepfer, D.D., Hauck, C.R. & Sieg, D.J. Signaling through focal adhesion kinase. Prog Biophys Mol Biol 71, 435–478 (1999).

8. Fluck, M., Carson, J.A., Gordon, S.E., Ziemiecki, A. & Booth, F.W. Focal adhesion proteins FAK and paxillin increase in hypertrophied skeletal muscle. Am J Physiol 277, C152–162 (1999).

9. Klossner, S., Durieux, A.C., Freyssenet, D. & Flueck, M. Mechano-transduction to muscle protein synthesis is modulated by FAK. European journal of applied physiology 106, 389–398 (2009).

10. Syeda, R., et al. LRRC8 Proteins Form Volume-Regulated Anion Channels that Sense Ionic Strength. Cell 164, 499–511 (2016).

11. Voss, F.K., et al. Identification of LRRC8 heteromers as an essential component of the volume-regulated anion channel VRAC. Science 344, 634–638 (2014).

12. Qiu, Z., et al. SWELL1, a Plasma Membrane Protein, Is an Essential Component of Volume-Regulated Anion Channel. Cell 157, 447–458 (2014).

13. Osei-Owusu, J., Yang, J., Vitery, M.D.C. & Qiu, Z. Molecular Biology and Physiology of Volume-Regulated Anion Channel (VRAC). Curr Top Membr 81, 177–203 (2018).

14. Sawada, A., et al. A congenital mutation of the novel gene LRRC8 causes agammaglobulinemia in humans. J Clin Invest 112, 1707–1713 (2003).

15. Kubota, K., et al. LRRC8 involved in B cell development belongs to a novel family of leucine-rich repeat proteins. FEBS Lett 564, 147–152 (2004).

16. Kumar, L., et al. Leucine-rich repeat containing 8A (LRRC8A) is essential for T lymphocyte development and function. J Exp Med 211, 929–942 (2014).

17. Abascal, F. & Zardoya, R. LRRC8 proteins share a common ancestor with pannexins, and may form hexameric channels involved in cell-cell communication. Bioessays 34, 551–560 (2012).

18. Cahalan, M.D. & Lewis, R.S. Role of potassium and chloride channels in volume regulation by T lymphocytes. Soc Gen Physiol Ser 43, 281–301 (1988).

19. Hazama, A. & Okada, Y. Ca2+ sensitivity of volume-regulatory K+ and Cl-channels in cultured human epithelial cells. J Physiol 402, 687–702 (1988).

20. Pedersen, S.F., Klausen, T.K. & Nilius, B. The identification of a volume-regulated anion channel: an amazing Odyssey. Acta Physiol (Oxf) 213, 868–881 (2015).

21. Pedersen, S.F., Okada, Y. & Nilius, B. Biophysics and Physiology of the Volume-Regulated Anion Channel (VRAC)/Volume-Sensitive Outwardly Rectifying Anion Channel (VSOR). Pflugers Arch 468, 371–383 (2016).

22. Eggermont, J., Trouet, D., Carton, I. & Nilius, B. Cellular function and control of volume-regulated anion channels. Cell Biochem Biophys 35, 263–274 (2001).

23. Zhang, Y., et al. SWELL1 is a regulator of adipocyte size, insulin signalling and glucose homeostasis. Nature cell biology 19, 504–517 (2017).

24. Gunasekar, S.K., Xie, L. & Sah, R. SWELL signalling in adipocytes: can fat ‘feel’ fat? Adipocyte 8, 223–228 (2019).

25. Browe, D.M. & Baumgarten, C.M. Stretch of beta 1 integrin activates an outwardly rectifying chloride current via FAK and Src in rabbit ventricular myocytes. J Gen Physiol 122, 689–702 (2003).

26. Browe, D.M. & Baumgarten, C.M. EGFR kinase regulates volume-sensitive chloride current elicited by integrin stretch via PI-3K and NADPH oxidase in ventricular myocytes. J Gen Physiol 127, 237–251 (2006).

27. Bruning, J.C., et al. A muscle-specific insulin receptor knockout exhibits features of the metabolic syndrome of NIDDM without altering glucose tolerance. Mol Cell 2, 559–569 (1998).

28. Moller, D.E., et al. Transgenic mice with muscle-specific insulin resistance develop increased adiposity, impaired glucose tolerance, and dyslipidemia. Endocrinology 137, 2397–2405 (1996).

29. Kim, J.K., et al. Redistribution of substrates to adipose tissue promotes obesity in mice with selective insulin resistance in muscle. J Clin Invest 105, 1791–1797 (2000).

30. Kang, C., et al. SWELL1 is a glucose sensor regulating beta-cell excitability and systemic glycaemia. Nat Commun 9, 367 (2018).

31. Haralampieva, D., et al. Human Muscle Precursor Cells Overexpressing PGC-1alpha Enhance Early Skeletal Muscle Tissue Formation. Cell Transplant 26, 1103–1114 (2017).

32. Ruas, J.L., et al. A PGC-1alpha isoform induced by resistance training regulates skeletal muscle hypertrophy. Cell 151, 1319–1331 (2012).

33. Singh, J., Verma, N.K., Kansagra, S.M., Kate, B.N. & Dey, C.S. Altered PPARgamma expression inhibits myogenic differentiation in C2C12 skeletal muscle cells. Molecular and cellular biochemistry 294, 163–171 (2007).

34. Conejo, R., Valverde, A.M., Benito, M. & Lorenzo, M. Insulin produces myogenesis in C2C12 myoblasts by induction of NF-kappaB and downregulation of AP-1 activities. J Cell Physiol 186, 82–94 (2001).

35. Lee, S. & Dong, H.H. FoxO integration of insulin signaling with glucose and lipid metabolism. J Endocrinol 233, R67–R79 (2017).

36. Gross, D.N., van den Heuvel, A.P. & Birnbaum, M.J. The role of FoxO in the regulation of metabolism. Oncogene 27, 2320–2336 (2008).

37. Barakat, A.I., Leaver, E.V., Pappone, P.A. & Davies, P.F. A flow-activated chloride-selective membrane current in vascular endothelial cells. Circ Res 85, 820–828 (1999).

38. Nakao, M., Ono, K., Fujisawa, S. & Iijima, T. Mechanical stress-induced Ca2+ entry and Cl-current in cultured human aortic endothelial cells. Am J Physiol 276, C238–249 (1999).

39. Nilius, B. & Droogmans, G. Ion channels and their functional role in vascular endothelium. Physiol Rev 81, 1415–1459 (2001).

40. Romanenko, V.G., Davies, P.F. & Levitan, I. Dual effect of fluid shear stress on volume-regulated anion current in bovine aortic endothelial cells. Am J Physiol Cell Physiol 282, C708–718 (2002).

41. Strange, K., Yamada, T. & Denton, J.S. A 30-year journey from volume-regulated anion currents to molecular structure of the LRRC8 channel. J Gen Physiol (2019).

42. Ma, Y., Fu, S., Lu, L. & Wang, X. Role of androgen receptor on cyclic mechanical stretch-regulated proliferation of C2C12 myoblasts and its upstream signals: IGF-1-mediated PI3K/Akt and MAPKs pathways. Mol Cell Endocrinol 450, 83–93 (2017).

43. Fu, S., Yin, L., Lin, X., Lu, J. & Wang, X. Effects of Cyclic Mechanical Stretch on the Proliferation of L6 Myoblasts and Its Mechanisms: PI3K/Akt and MAPK Signal Pathways Regulated by IGF-1 Receptor. Int J Mol Sci 19 (2018).

44. Shan, X., et al. Suppression of Grb2 expression improved hepatic steatosis, oxidative stress, and apoptosis induced by palmitic acid in vitro partly through insulin signaling alteration. In Vitro Cell Dev Biol Anim 49, 576–582 (2013).

45. Liu, X., et al. Downregulation of Grb2 contributes to the insulin-sensitizing effect of calorie restriction. American journal of physiology. Endocrinology and metabolism 296, E1067–1075 (2009).

46. Mitra, P. & Thanabalu, T. Myogenic differentiation depends on the interplay of Grb2 and N-WASP. Biochim Biophys Acta Mol Cell Res 1864, 487–497 (2017).

47. Affourtit, C. Mitochondrial involvement in skeletal muscle insulin resistance: A case of imbalanced bioenergetics. Biochimica et biophysica acta 1857, 1678–1693 (2016).

48. Seale, P., et al. PRDM16 controls a brown fat/skeletal muscle switch. Nature 454, 961–967 (2008).

49. Bothe, G.W., Haspel, J.A., Smith, C.L., Wiener, H.H. & Burden, S.J. Selective expression of Cre recombinase in skeletal muscle fibers. Genesis 26, 165–166 (2000).

50. Rotwein, P. & Wilson, E.M. Distinct actions of Akt1 and Akt2 in skeletal muscle differentiation. J Cell Physiol 219, 503–511 (2009).

51. Heron-Milhavet, L., Mamaeva, D., Rochat, A., Lamb, N.J. & Fernandez, A. Akt2 is implicated in skeletal muscle differentiation and specifically binds Prohibitin2/REA. J Cell Physiol 214, 158–165 (2008).

52. Chen, L., Becker, T.M., Koch, U. & Stauber, T. The LRRC8/VRAC anion channel facilitates myogenic differentiation of murine myoblasts by promoting membrane hyperpolarization. J Biol Chem (2019).

53. Chen, X. & Chalfie, M. Modulation of C. elegans touch sensitivity is integrated at multiple levels. The Journal of neuroscience : the official journal of the Society for Neuroscience 34, 6522–6536 (2014).

54. Kim, J., Bilder, D. & Neufeld, T.P. Mechanical stress regulates insulin sensitivity through integrin-dependent control of insulin receptor localization. Genes Dev 32, 156–164 (2018).

55. Relaix, F., et al. Pax3 and Pax7 have distinct and overlapping functions in adult muscle progenitor cells. The Journal of cell biology 172, 91–102 (2006).

56. Buckingham, M., et al. Myogenic progenitor cells in the mouse embryo are marked by the expression of Pax3/7 genes that regulate their survival and myogenic potential. Anat Embryol (Berl) 211 Suppl 1, 51–56 (2006).

57. Ostler, J.E., et al. Effects of insulin resistance on skeletal muscle growth and exercise capacity in type 2 diabetic mouse models. American journal of physiology. Endocrinology and metabolism 306, E592–605 (2014).

58. Reusch, J.E., Bridenstine, M. & Regensteiner, J.G. Type 2 diabetes mellitus and exercise impairment. Rev Endocr Metab Disord 14, 77–86 (2013).

59. Jaiswal, N., et al. The role of skeletal muscle Akt in the regulation of muscle mass and glucose homeostasis. Mol Metab 28, 1–13 (2019).

## References

1. Zhang, Y., et al. SWELL1 is a regulator of adipocyte size, insulin signalling and glucose homeostasis. Nature cell biology 19, 504–517 (2017).

2. Cong, L., et al. Multiplex genome engineering using CRISPR/Cas systems. Science 339, 819–823 (2013).

3. Kang, C., et al. SWELL1 is a glucose sensor regulating beta-cell excitability and systemic glycaemia. Nat Commun 9, 367 (2018).

4. Hindi, L., McMillan, J.D., Afroze, D., Hindi, S.M. & Kumar, A. Isolation, Culturing, and Differentiation of Primary Myoblasts from Skeletal Muscle of Adult Mice. Bio Protoc 7 (2017).

5. Bonetto, A., Andersson, D.C. & Waning, D.L. Assessment of muscle mass and strength in mice. Bonekey Rep 4, 732 (2015).

6. Dougherty, J.P., Springer, D.A. & Gershengorn, M.C. The Treadmill Fatigue Test: A Simple, High-throughput Assay of Fatigue-like Behavior for the Mouse. Journal of visualized experiments : JoVE (2016).

7. Wende, A.R., et al. Enhanced cardiac Akt/protein kinase B signaling contributes to pathological cardiac hypertrophy in part by impairing mitochondrial function via transcriptional repression of mitochondrion-targeted nuclear genes. Molecular and cellular biology 35, 831–846 (2015).

